# Generative modeling of biological shapes and images using a probabilistic *α*-shape sampler

**DOI:** 10.1101/2024.01.09.574919

**Authors:** Emily T. Winn-Nuñez, Hadley Witt, Dhananjay Bhaskar, Ryan Y. Huang, Jonathan S. Reichner, Ian Y. Wong, Lorin Crawford

**Affiliations:** Division of Applied Mathematics, Brown University, Providence, RI, USA; Graduate Program in Pathobiology, Brown University, Providence, RI, USA; Division of Surgical Research, Department of Surgery, Rhode Island Hospital, Providence, RI, USA; Department of Genetics, Yale School of Medicine, New Haven, CT USA; Department of Computer Science, Brown University, Providence, RI USA; School of Engineering, Legoretta Cancer Center, Brown University, Providence, RI USA; Microsoft Research, Cambridge, MA, USA; Department of Biostatistics, Brown University, Providence, RI, USA; Center for Computational Molecular Biology, Brown University, Providence, RI, USA

## Abstract

Understanding morphological variation is an important task in many areas of computational biology. Recent studies have focused on developing computational tools for the task of sub-image selection which aims at identifying structural features that best describe the variation between classes of shapes. A major part in assessing the utility of these approaches is to demonstrate their performance on both simulated and real datasets. However, when creating a model for shape statistics, real data can be difficult to access and the sample sizes for these data are often small due to them being expensive to collect. Meanwhile, the current landscape of generative models for shapes has been mostly limited to approaches that use black-box inference—making it difficult to systematically assess the power and calibration of sub-image models. In this paper, we introduce the *α*-shape sampler: a probabilistic framework for generating realistic 2D and 3D shapes based on probability distributions which can be learned from real data. We demonstrate our framework using proof-of-concept examples and in two real applications in biology where we generate (*i*) 2D images of healthy and septic neutrophils and (*ii*) 3D computed tomography (CT) scans of primate mandibular molars. The *α*-shape sampler R package is open-source and can be downloaded at https://github.com/lcrawlab/ashapesampler.

**Author Summary:** Using shapes and images to understand genotypic and phenotypic variation has proven to be an effective strategy in many biological applications. Unfortunately, shape data can be expensive to collect and, as a result, sample sizes for analyses are often small. Despite methodological advancements in shape statistics and machine learning, benchmarking standards for evaluating new computational tools via data simulation is still underdeveloped. In this paper, we present a probability-based pipeline called the *α*-shape sampler which has the flexibility to generate new and unobserved shapes based on an input set of data. We extensively evaluate the generative capabilities of our pipeline using 2D cellular images of neutrophils and 3D mandibular molars from two different suborders of primates.

## Introduction

Shape statistics has become an integral component of several applications within computational biology including medical imaging ^1^, geometric morphometrics ^2–4^, and cell biology ^5,6^. Recently, there has been a focus to develop computational tools that address the subimage analysis problem: given a collection of images or shapes, find the features that best explain the variation between them with respect to a response variable ^7^. One example of this type of analysis is identifying the biologically-relevant atomic and residue-level differences between two protein structural ensembles ^8^. To date, several approaches have been proposed with the aim to quantify the global variation between images and shapes including some in topological data analysis ^9–12^, methods leveraging landmark-based ^13–15^ or diffeomorphic-based representations ^2,16–18^, and tools that use “functional maps” to identify similarities and differences between shapes via a learned latent space ^19^.

Despite the many methodological advances being made for the subimage selection problem in shape analysis, there has yet to be a principled framework to assess the power and limitations of these new tools. Traditionally, there are two common strategies for benchmarking feature selection methods in computational biology: (i) by analyzing real data where there is a “ground truth” about which features are associated with a given phenotype of interest, or (ii) by using simulations where synthetic data is generated such that we know the causal relationship between features and the response variable. Both of these strategies have well-established statistical practices for tabular data (e.g., gene expression in genomics) but they become increasingly difficult to implement when working with shapes. Using data from real biomedical studies for methodological benchmarking is a challenge because shape-based modalities can be hard to collect. On the other hand, when data is able to be collected, sample sizes within studies are usually small, which both compromises the statistical power of the methods being assessed and inhibits the ability to study algorithmic robustness to variance between observations. Lastly, the relationship between shape and phenotype is largely speculative for many biological applications. For example, there have been radiomic studies which have proposed an association between tumor morphology and survival prognostics for patients with glioblastoma, but the exact biological mechanisms connecting the two remains unknown ^1,20^.

Simulation studies are an alternative way to evaluate newly developed computational tools in shape analyses. The key to performing these studies is to have an interpretable generative model such that the process for creating synthetic (yet realistic) shapes is well understood. This facilitates the ability to assess how powered a tool is at identifying causal features driving the morphological variation across samples. In general, algorithmic frameworks for generating synthetic shapes consists of two steps: (i) a procedure to generate a point set and (ii) a set of rules for reconstructing a shape from those points. Multiple end-to-end shape generation pipelines have been introduced in the literature but each have their own sets of limitations. For example, to sample random points from a probability distribution over a manifold, one theoretically needs to know the manifold itself which can be impractical to estimate for many applications ^21–24^. Recently, there are have been machine learning algorithms that have been developed for generating point clouds and reconstructing shapes using dual generators ^25^, diffusion-based methods ^26^, encoders ^27^, and generative adversarial networks ^28,29^; but each of these frameworks lack transparency into the generative process for creating new synthetic shapes ^30^. From a more mathematical perspective, several methods have been proposed to infer shapes from randomly generated point clouds. Many of these approaches use C^̌^ech and Vietoris Rips complexes ^31,32^; however, unfortunately, they require tens (and sometimes hundreds) of simplicial complexes to be constructed for each point set resulting in long runtimes. There are 2D shape reconstruction methods based on contours ^33^ and curves ^34^, but their theory does not directly translate to higher dimensional objects ^35^. Lastly, probability-based shape generative pipelines are still in their infancy and have thus far relied on component vector analysis where parts of 2D and 3D objects are broken into smaller components and the assembly/connectivity between components are hidden variables learned by a pre-specified model ^36,37^.

In this work, we present the *α*-shape sampler: a probabilistic framework which takes in a collection of real shapes or images as input and generates new synthetic ones with features that both quantitatively and qualitatively resemble data in the input set. Methodologically, *α*-shapes require a single numerical parameter *α* for reconstruction which can be interpreted as a measure of shape detail or granularity (Fig 1). They can also be generated in *O*(*P* log *P*) time where *P* is the number of points in the point cloud that is input into the algorithm ^38^. As part of our contributions, we introduce a scalable näıve, data-driven algorithm to estimate the *reach* ^39^ for a given set of shapes and theoretically relate it to the numerical *α* parameter. Altogether these properties allow our proposed framework to scale and accommodate the growing sizes of emerging imaging and shape-based databases. It is worth mentioning that, while the mathematical concept of reach has been used extensively in topological data analysis to reconstruct shapes and sample point clouds ^40,41^, to our knowledge, we are the first to tie it *α*-shapes parameter for an end-to-end generative modeling pipeline. Shape generation using *α*-shapes has been previously studied in two-dimensions where the underlying manifold is known ^42^ and to learn about shape boundaries ^43,44^; while shape reconstruction with *α*-shapes has primarily been studied in three dimensions ^45–47^. They have also been previously used structural biology application in ecology^48,49^ but, overall, the focus of these studies was to understand the interpretation of the parameter of *α* itself rather than attempting to use *α*-shapes as a basis to create a framework for generating new data.

**Figure 1.**
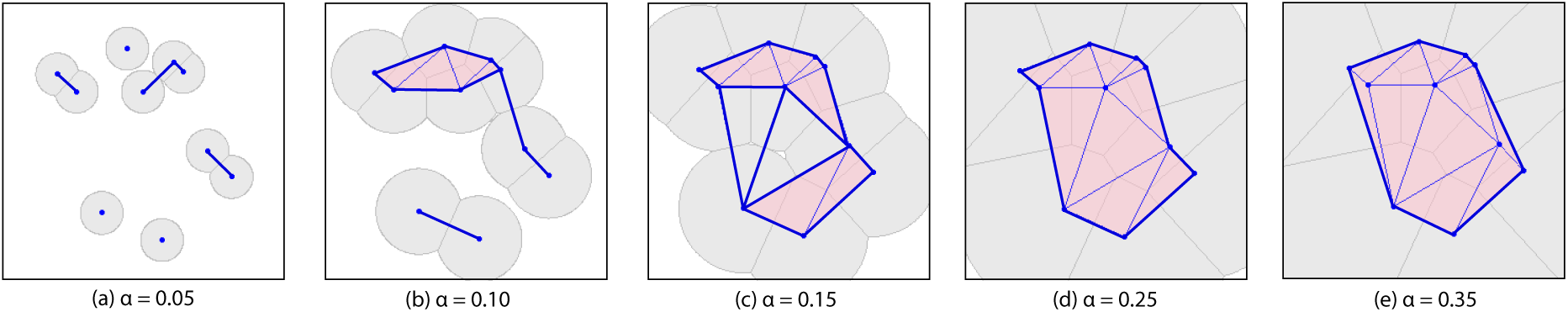
An example of various *α*-shapes for the same set of points under different choices for the numerical parameter *α*. Here, we consider different parameter values (a) *α* = 0.05, **(b)**, *α* = 0.10, (c) *α* = 0.15, (d) *α* = 0.2, and (e) *α* = 0.35. In each panel, the gray shapes are the intersection of balls of radius *α* and the Voronoi cells at each point. The pink triangles are then faces representing the collective interior, and the blue lines are edges of the *α*-complex. The bold blue edges are known as the “boundary edges” and denote the *α*-shape for each panel. In **(a)** and **(b)**, where *α* is smaller, we have disconnected components. In **(c)**, we see an instance where edges may form the boundary of a face, but the face is not quite yet filled in since the three Voronoi cells have not collectively met. In **(d)**, the faces are filled in and one of the points becomes an interior point while the rest remain *α*-extreme points. In **(e)**, *α* is large enough such that the given *α*-complex is the Deluanay triangulation and convex hull of the point set. When determining how to generate a new shape from an existing dataset, we use information within the given simplicial complex to determine how many points are needed, where the points should be sampled, and the appropriate *α* parameter to connect the points. For a more detailed overview and theoretical discussion of concepts surrounding *α*-shapes, see Materials and Methods and Supporting Information.

Throughout the rest of the paper, we will describe the *α*-shape sampler using a combination of probability theory, topology, and tools from differential geometry. We then translate the theoretical components of the pipeline into a series of algorithmic steps for practical implementation. Finally, we illustrate the utility of our approach on small proof-of-concept examples (annuli in two dimensions and tori in three dimensions) and real datasets (neutrophils in two dimensions and primate mandibular molars in three dimensions). We find that the *α*-shape sampler is effective at generating new shapes which honor major local and global characteristics of realistic data, while also maintaining algorithmic transparency so that the pipeline can be used for a wide-range of biological applications.

## Results

### Algorithmic overview of the *α*-shape sampler

Statistically, *α*-shapes are convenient because they require a single numerical parameter *α* to encode all connectivity information for a point set. For example, in Fig 1a-c, we see that all points are *α*-extreme (i.e., on the border); while in Fig 1d, we see that *α* becomes large enough such that one point is not *α*-extreme and is therefore an *interior* point of the shape. Finally, in Fig 1e, there are three interior points and the rest are boundary or *α*-extreme points. An extension of this figure showing different *α*-shapes being formed as a function of the number of points sampled from a unit square and the parameter *α* can be found in Fig S1. With this theory in mind, a probability distribution on *α*-shapes can be explicitly estimated via uniform point sampling on a given (approximate) manifold and then shapes can be constructed from that point set using *α* (see Supporting Information). Recent work has investigated using the *α* parameter as a shape characteristic ^48,49^ but, to our knowledge, it has yet to be used for shape generation. This is likely due to the requirement that point sets need to be in general position, a characteristic often not seen in nature. However, we work within the confines of this assumption in return for theoretical soundness, statistical simplicity, and algorithmic transparency.

We will detail our probabilistic generative framework while assuming that we are working with shapes that are *d* = 2 or 3-dimensions. The *α*-shape sampler involves five key steps (see Fig 2a). To begin, the pipeline receives real shapes; throughout the rest of this paper, we will refer to these input data as “reference” shapes. Note that we depict these reference shapes as binary masks in Fig 2, but the *α*-shape sampler software can take shape data in any format as input. In the second step, the reference shapes are aligned, scaled (if applicable), and converted to triangular meshes which we treat as simplicial complexes. In the third step, the reference meshes are used in a generative algorithm which, in the fourth step, outputs newly generated shapes in the form of new *α*-complexes. In the fifth and final step, these newly generated *α*-complexes are converted back into binary masks (or any other data representation), to match the same format as the original input reference data.

**Figure 2.**
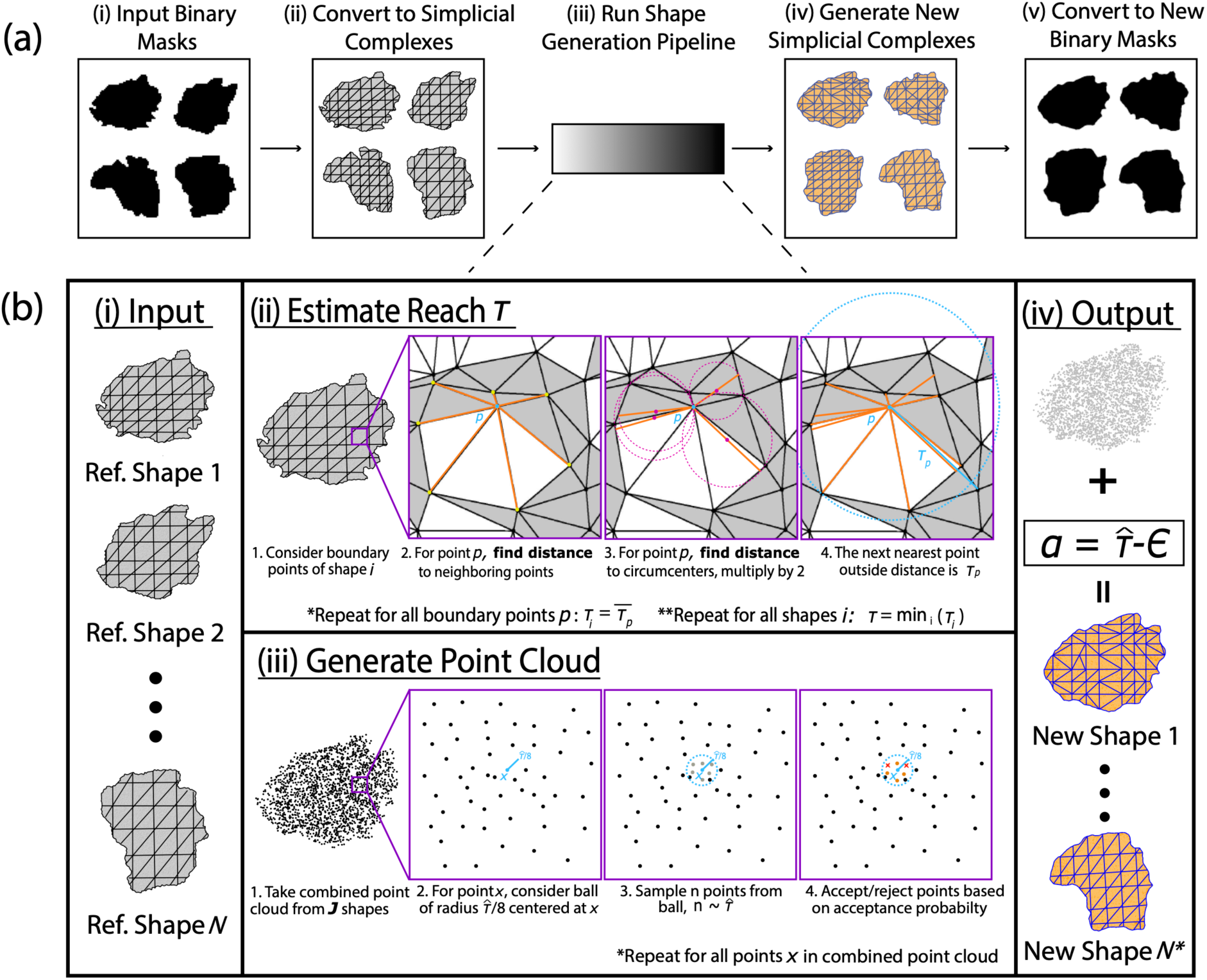
Schematic overview of the *α*-shape sampler: a probabilistic framework for simulating realistic 2D and 3D images and shapes. (a) A general illustration of the pre- and post-processing workflow in the *α*-shape sampler software. In step (i), the user inputs data of real shapes in some format—in this case, binary masks for illustration. We refer to these data as “reference” shapes. In step (ii), the reference masks are converted to triangular meshes which are treated as simplicial complexes. In step (iii), the reference meshes are input into the shape generation pipeline which, in step (iv), outputs newly generated shapes in the form of *α*-complexes. Finally, in step (v), these generated *α*-complexes are converted back to match the same format as the original input data (again, here, binary masks). **(b)** Details underlying the algorithm for generating new shapes via the *α*-shape sampler. (i) A collection meshes from *N* reference shapes are given to the software. For simplicity, we assume that these shapes are from the same phenotypic class and, thus, their points are from the same manifold. (ii) Next, we estimate the reach *τ_i_* for each reference shape by computing the distance to edge neighbors for each point (i.e., vertex in the mesh) and the circumcenters to neighboring faces (note that we also evaluate tetrahedra for 3D objects). The next closest vertex is the value *τ_p_* for point *p*, and the smallest *τ_p_* among all points is the value of *τ_i_* for the *i*-th reference shape. We then take the minimum *τ* = (*τ*_1_*, …, τ_N_*) to be the representative estimate of the reach 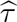 for all reference shapes. (iii) We create a partial point cloud by combining points from *J* reference shapes in our input dataset, where 2 *≤ J ≤ N*. Next, we sample new points from a ball of radius 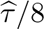 around vertices in the partial point cloud. Each new point is accepted or rejected according to a probability-based rule. (iv) Once we have the newly sampled point cloud, we set 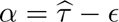, where *ε >* 0 is arbitrarily small, and generate the *α*-complexes for new shapes.

We assume that all reference shapes from a phenotypic class (e.g., healthy cells or molars from a given species of primate) have vertices sampled from the same underlying manifold and that the variation observed across shapes within the class stems from a finite sampling of points. With this in mind, the generative algorithm proportion of the *α*-shape sampler is comprised of four main steps (see Fig 2b).

First, the *N* collection reference meshes are input into the algorithm. We represent the *i*-th reference mesh as *K_i_*= *{V_i_, E_i_, F_i_, T_i_}* which is collection of vertices *V_i_*, edges *E_i_*, faces *F_i_*, and tetrahedra *T_i_* (if applicable). In the second step, we estimate the reach *τ_i_* for every *i*-th reference mesh by computing the distance to edge neighbors and the circumcenter distance to neighboring faces (and tetrahedra for 3D objects) for each boundary vertex in the complex *p ∈ ∂K_i_*. After completing this for all *N* reference shapes, we have a vector of shape-specific reach estimates ***τ*** = (*τ*_1_*, …, τ_N_*). In the third step, we select 2 *≤ J ≤ N* reference shapes from the input dataset which we use as a basis to generate new shapes. Here, we combine the point clouds from the *J* shapes into a joint partial point and take the minimum between their corresponding values in ***τ*** to be the reach estimate 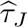. Next, we sample candidate points for the newly generated shapes from balls of radius 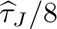 around vertices in the joint partial point cloud. A radius of 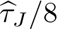 is chosen to force newly sampled points to remain relatively close to the boundary of the reference point cloud. Each new candidate point is accepted or rejected according to a probability-based rule with parameter *θ* (see Materials and Methods). The *θ* parameter is the minimum number of points in the joint partial point cloud that need to neighbor the new candidate point in order to accept it. It effectively determines the level of confidence needed to believe that a randomly sampled point is from the same underlying manifold as the reference data. Once we have the newly sampled point cloud, in the fourth step of the algorithm, we set 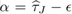, where *ε >* 0 is arbitrarily small, and generate the *α*-complexes for new shapes. By default, the *α*-shape sampler software sets *J* = 2, *θ* = *d* (i.e., same dimensions as the input data), and *ε* = 0.001 (see URLs). Unless otherwise stated, these are the values that we also use to generate all of the results presented throughout the rest of the paper.

There are two important components to the implementation of our pipeline. First, the *α*-shape sampler uses a function to compute the reach for each shape that is completely separate from the shape generation function (again see Fig 2b). This serves two purposes: (i) it increases computational speed by avoiding redundant calculations, and (ii) it provides an informal check for potential outlier shapes before using those shapes as reference inputs for the generative part of algorithm (e.g., this can be done by empirically assessing the tails of the distribution for ***τ***). Second, setting *α* to be just under 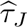 for some subset of reference shapes guarantees that we will preserve the original homology and most of the local geometry that is present in the reference dataset without losing any features or generating any atypical ones. Theoretical details of our implementation are fully detailed in the Materials and Methods and Supporting Information.

### 2D proof-of-concept study with simulated annuli

To demonstrate the *α*-shape sampler, we begin with a two-dimensional (2D) toy example where we simulate *N* = 50 “real” (i.e., reference) annuli with inner radius *r* = 0.25, outer radius *R* = 0.75, and thickness equal to *R − r* = 0.5. Each reference annulus is constructed by sampling *P* = 500 points uniformly from the annulus and then connecting them using true *α* = 0.15. The reach value for these reference annuli is given by the inner radius of the hole such that *τ* = 0.25. We consider these measurements to be the “ground truth” during evaluation.

Using the real 2D annuli as input data, we generate another *N ^∗^* = 10 annuli using the *α*-shape sampler. Figs 3a and 3b show that the generated annuli preserve the homology of the original reference shapes (i.e., each generated shape is singular connected component and has exactly one hole). To further evaluate how “realistic” the geometric characteristics were for the newly generated annuli, we first identified their *α*-extreme points and separated them into two categories: (i) radii less than 0.5 and (ii) radii greater than 0.5. The averages of both categories were used to numerically define each generated shape’s inner and outer radii, respectively. The thickness of each generated shape was then found by subtracting the inner radius from the outer radius. Table S1 gives the root mean square error (RMSE) for each of these characteristics when comparing the generated annuli to the real reference annuli. Overall, we see relatively low RMSEs (values below 0.01 for each category) which aligns with the aesthetic similarity between the shapes seen in Fig 3. It is important to note that the mean estimated reach for the generated annuli produced by the *α*-shape sampler was 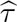 = 0.1749 *±* 0.009. While less than the true value of *τ* = 0.25, this is unexpected given that we are estimating the reach from the data directly (rather than estimating it using the true radius). Indeed, we would rather our estimate of the reach be smaller than the truth and, consequently, have to sample more points rather than our estimate of *τ* be too large and we lose geometric information about the shapes.

**Figure 3.**
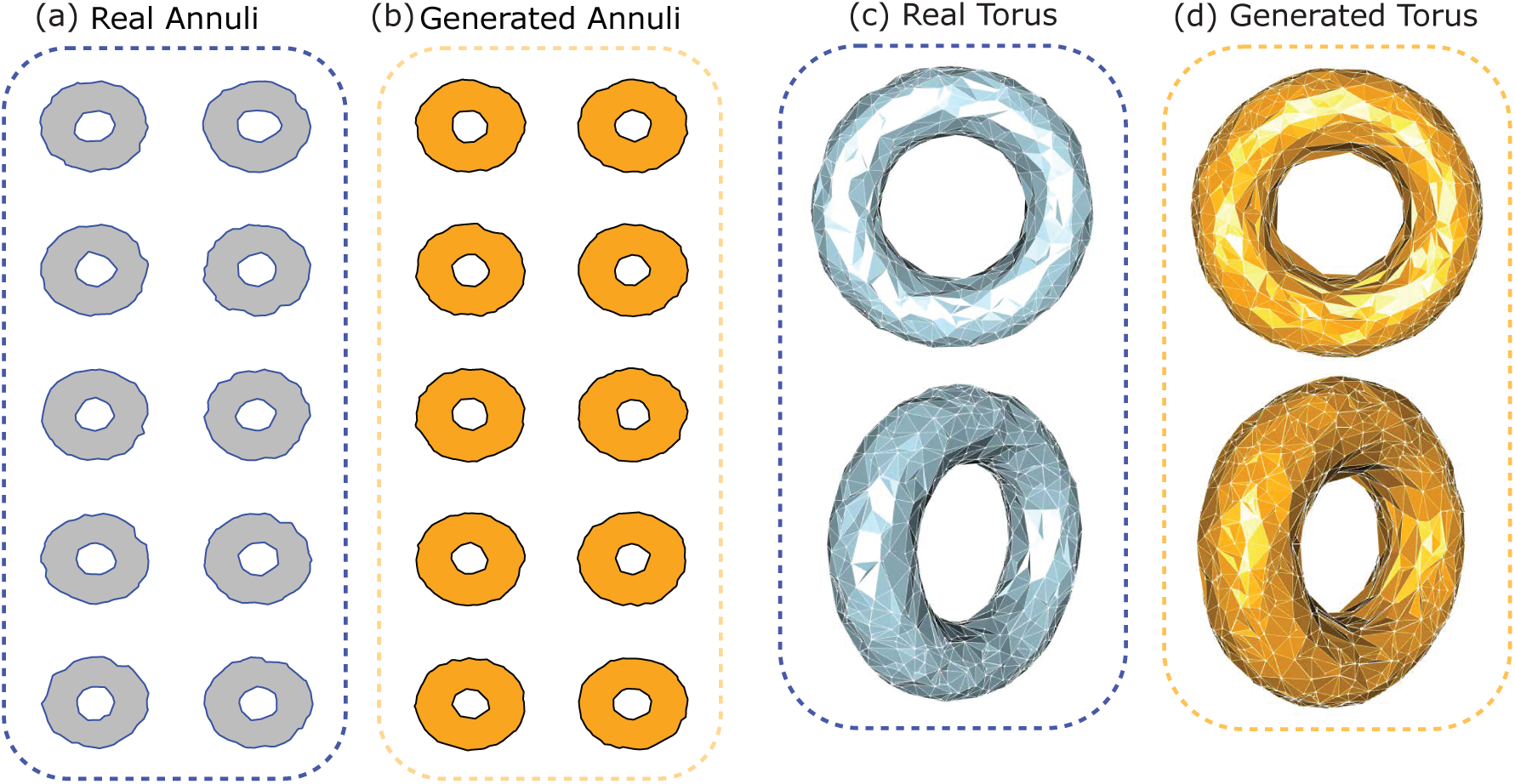
Qualitative comparisons of real and generated 2D annuli and 3D tori using the *α*-shape sampler. Panels **(a)** and **(b)** show real (gray) and generated (orange) annuli. Similarly, in panels **(c)** and **(d)**, we show real (gray) and generated (orange) tori. Overall, we see that the *α*-shape sampler generates slightly thicker shapes than the examples in the original dataset (see Tables S1 and S2 for a quantitative evaluation). Nonetheless, the generated shapes preserve the most important topological property in that they all have exactly one connected component and exactly one hole.

### 3D proof-of-concept study with simulated tori

Next, we extend our demonstration of the *α*-shape sampler to a three-dimensional (3D) toy example where we simulate *N* = 50 “real” (i.e., reference) tori with major radius *R* = 0.75 and minor radius *r* = 0.25. Each reference torus was constructed by sampling *P* = 5000 points uniformly using the alphashape3d R package ^50^ with the Computational Geometry Algorithms Library (CGAL) ^51^ where the points were connected using a true *α* = 0.25. The reach value for these reference tori is *τ* = 0.5 which corresponds to the radius of the hole (or tube) of the tori. As with the previous proof-of-concept study using the 2D annuli, we again consider the geometric measurements for the real reference tori to be the “ground truth” during our assessment. Examples of the real reference tori can be found in Fig 3c.

Using the real 3D tori as input data, we generate another *N ^∗^* = 10 tori using the *α*-shape sampler. An example of these generated shapes can be found in Fig 3d. Here, we see that the generated tori qualitatively preserve the homology of the original data where each have one connected component and one hole. To get estimates of the major and minor radii for the generated torus, we start by examining their boundary points. The following relates the major and minor radii for a torus centered at the origin

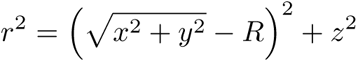

where (*x, y, z*) are the Cartesian coordinates of the boundary points for the torus. Rearranging the above equation then yields the following relationship

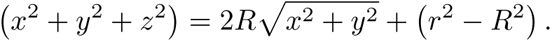

By treating *Y* = *x*^2^ + *y*^2^ + *z*^2^ to be a response variable, 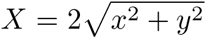 to be a covariate, *β* = *R* to be a coefficient, and *ε* = (*r*^2^ *− R*^2^) to be a residual, the above rewritten equation mirrors a linear model. As a result, we can use ordinary least squares to estimate the appropriate values for (*β, ε*). This then allows us to infer corresponding estimates for the major *R* and minor *r* radii for each generated torus, respectively.

Table S2 compares the major and minor radii estimates for the tori generated by the *α*-shape sampler to same characteristics in the original reference shapes. We see that while the major radius *R* is well preserved (RMSE = 0.002), the minor radius *r* is slightly larger for the generated shapes (RMSE = 0.02). This result translated to a slightly larger thickness for the generated tori (again see Fig 3d). While generally still close, it does demonstrate a potential shortcoming in our data-driven approach for shape generation where our random sampling algorithm can be prone to accept points outside of the reference boundary, particularly for shapes with smooth surfaces. While this issue may be corrected via some post-processing step to assure that the generated shapes are on a desired scale, we still caution that the probabilistic nature of the *α*-shape sampler is not perfect and may lead to a slight distortion of shape geometry. It is still worth noting that, despite the slightly larger thickness, the mean reach estimate for the generated tori produced by our algorithm was 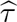 = 0.402 *±* 0.006, lower than the true value of *τ* = 0.5. Again, the lower reach estimate helps us to preserve the majority of geometric and topological characteristics in the generated shapes even if the overall scale is slightly misrepresented.

### Comparison of real and generated shapes based on primary human neutrophils from healthy and septic patients

Human cells display diverse and dynamic morphologies, driven by the rich interplay between the intracellular cytoskeleton and matrix adhesion during cell migration ^52,53^. For example, neutrophils are versatile “first responders” of the innate immune system that are rapidly recruited to tissue sites of injury and infection ^54^. Neutrophils become adherent and polarized after “activation” by proinflammatory mediators ^55^, exhibiting a leading edge with protrusive pseudopods as well as a trailing edge with a contractile uropod ^56^. Indeed, such polarized morphologies appear to be correlated with faster neutrophil motility, but can be considerably more heterogeneous for slower moving cells ^57^. Further, neutrophils exhibit pro-found defects in migration and antimicrobial function during sepsis, an aberrant host response to infection that can result in multi-organ failure and death ^58^. An unresolved problem is to meaningfully classify neutrophils, since they plastically transition through distinct phenotypic states but also occur as distinct subsets defined by biomarkers and gene expression ^59^.

As a first case study, we applied the *α*-shape sampler to two-dimensional cell shapes acquired from phase microscopy images of primary human neutrophils. Briefly, neutrophils were isolated from consented healthy donors and septic patients at Rhode Island Hospital (with approval from the Institutional Review Board), then plated at compliant polyacylamide hydrogel substrates functionalized with fibronectin (see Materials and methods and Witt et al. ^60^ for more details). Representative cell morphologies were manually traced, converted to binary masks, and then turned into simplicial complexes (similar to what was shown in Fig 2a). The *α*-shape sampler was used to synthetically generate additional cells using the default parameters *J* = 2, *θ* = 2, and ε = 0.001. The training set consisted of approximately *N* = 20 neutrophil shapes each from the healthy donors and septic patients, which were then used to generate *N ^∗^* = 25 new neutrophil shapes from each class. Qualitatively, real healthy neutrophils exhibited relatively rounded and compact morphologies (including a uropod ^56^) with a typical diameter of *∼*10 microns (*µ*m), which were visually similar to the generated healthy neutrophils (Fig 4a). In comparison, real septic neutrophils exhibited greater ruffling and elongated protrusions relative to real healthy neutrophils, which was visually recapitulated in the generated septic neutrophils (again see Fig 4a). Further, septic neutrophils showed greater spread areas than healthy neutrophils, with diameters approaching 15-20 *µ*m. These differences between healthy and septic neutrophil shapes were captured in the reach estimates produced by the *α*-shape sampler, with the healthy neutrophils having mean 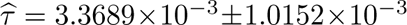 compared to the septic neutrophils having mean 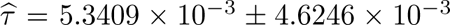. The larger mean *τ* can be explained by the larger variation along the border of the septic neutrophils, while the larger standard deviation reflects the greater single cell heterogeneity in shape.

**Figure 4.**
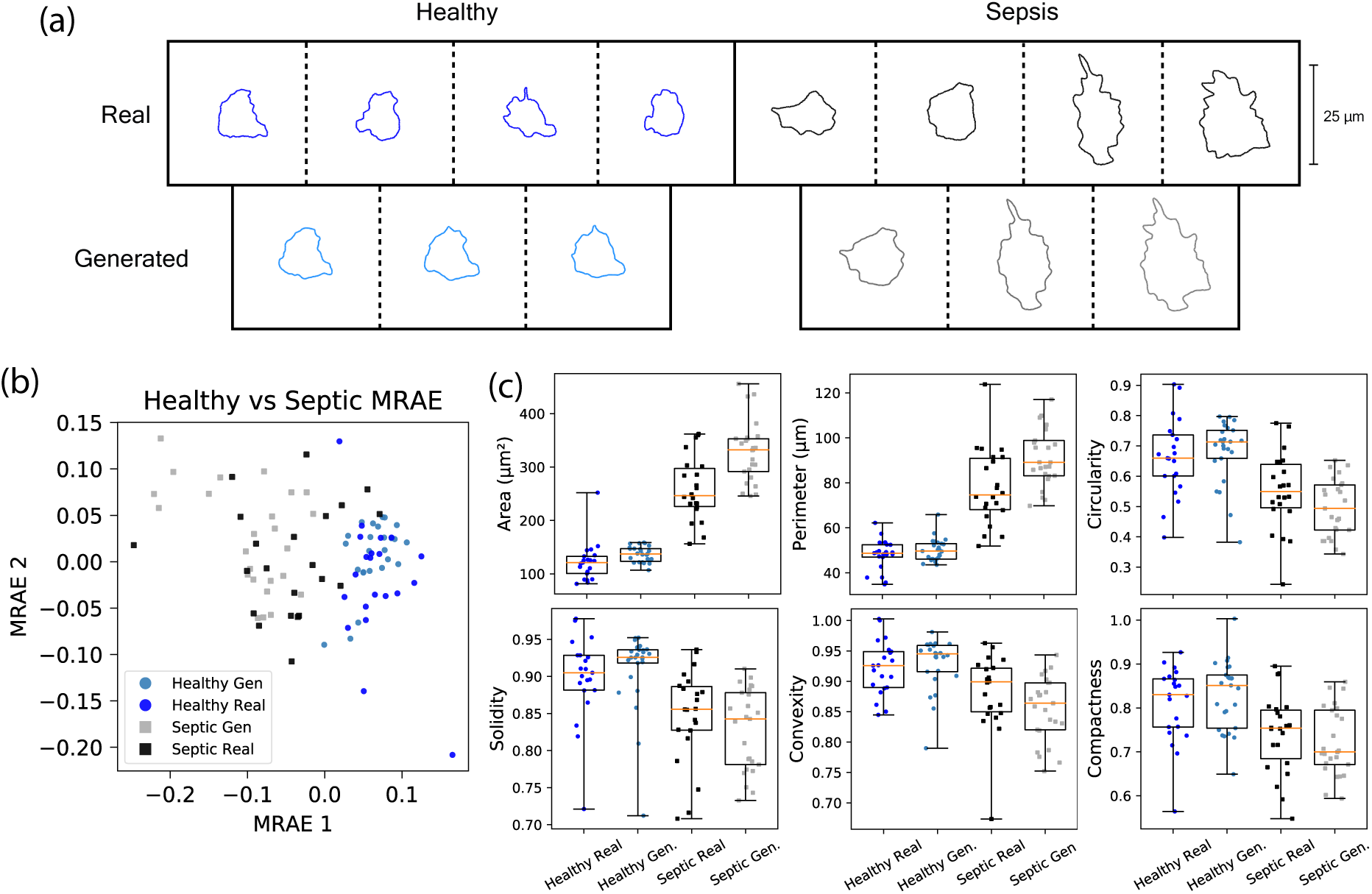
Application of the *α*-shape sampler to generate synthetic 2D images of healthy and septic neutrophils. **(a)** Examples of real healthy (blue), generated healthy (light blue), real septic (black), and generated septic (gray) neutrophils in gels with stiffness 1.5 kilopascals (kPa). Each synthetic neutrophil in the second row was generated using the two shapes it sits in between in the row above. Variation in the newly generate cells can be most seen along the boundary, which is a function of the sampling process in the *α*-shape pipeline. When comparing the generated and real cells, perhaps most noticeable are (i) the differences in area and (ii) the number of protrusions in the healthy versus septic cells. **(b)** We use a manifold regularized autoencoder (MRAE) to show that the generated shapes cluster and intermix with real cells in their respective categories. This provides evidence that the images being generated by the ***α***-shape sampler are realistic. **(c)** We compute the area, perimeter, circularity, solidity, convexity, and compactness of each real and generated cell. Next, we compare the distribution of these measurements for the healthy and septic groups, respectively. Here, if the *α*-shape is able to preserve geometric and morphological characteristics while generating new data, then we would expect the distributions of these measurements to line up within a group. Note that due to the high heterogeneity and difficulty aligning shapes, the generated septic neutrophils are slightly larger in area and perimeter than the real ones. However, the generated neutrophils with the *α*-shape sampler still capture other key shape characteristics.

To further quantify the differences between the real healthy and septic neutrophils and the similarities between real and generated neutrophils, 33 shape characteristics were calculated including area, perimeter length, compactness, and number of protrusions (see Materials and methods and Bhaskar et al. ^61^ for more details). These vectors were then projected onto a two-dimensional space using a manifold regularized autoncoder (MRAE) ^62^ as applied to Potential of Heat-diffusion for Affinity-based Transition Embedding (PHATE) coordinates (Fig 4b). In this lower dimensional representation, real healthy neutrophils are roughly grouped together for larger MRAE1, while real septic neutrophils are roughly grouped together for smaller MRAE1; although, there is not a large separation of these two groupings. Moreover, generated healthy neutrophils also group together with real healthy neutrophils for larger MRAE1, while generated septic neutrophils group together with real septic neutrophils for smaller MRAE1. These general trends were confirmed to be independent of the choice of dimension reduction method, including Uniform Manifold Approximation (UMAP) ^63^, PHATE ^64^, Principal Component Analysis (PCA), and a generalized autoencoder with an Adam optimizer and mean square error loss (Fig S3). For PCA, in particular, the top two principal components were most heavily weighted by area and perimeter in the loadings. Although MRAE is more difficult to interpret due to the nonlinear representation, the components were similarly weighted by area and perimeter but also solidity and circularity (based on inspection of cell shapes for varying MRAE1 and MRAE2).

Additional examination of these shape metrics revealed statistically significant quantitative differences between healthy and septic neutrophils (Fig 4c and Table S3). Notably, healthy real and generated neutrophils had comparable median area of *∼*125 *µ*m^2^ (*P* -value = 0.066). Moreover, septic real neutrophils had a median area of 246 *µ*m^2^, but septic generated neutrophils had a significantly larger median area of 332 *µ*m^2^ (*P* -value = 1.71 *×* 10*^−^*^4^). Similarly, healthy real and generated neutrophils had median perimeters of *∼*47 *µ*m (*P* -value = 0.189), while septic real neutrophils had a median perimeter of 75 *µ*m and septic generated neutrophils had a median perimeter of 89 *µ*m (*P* -value = 0.0032). In comparison, circularity (expressed as a ratio between 0 and 1 describing similarity to a circle, with 1 denoting a perfect circle), solidity (the fraction of the area of the cell over the area of the convex hull), convexity (the ratio of the convex hull perimeter to the cell perimeter), and compactness (the ratio of the diameter of the circle with the same area of the cell to the major axis of rectangular fit) showed statistically significant differences between the real healthy and septic neutrophils that were maintained by the generated healthy and septic neutrophils, but no statistically significant differences between the real and generated neutrophils. In order to elucidate this discrepancy between real and generated septic neutrophil shapes, we reexamined how the *α*-shape generator was sampling from the training set to define a “manifold” based on the union of point clouds from *J* = 2 reference shapes (see Material and methods). Without perfect alignment, in this setting, the corresponding combined manifold will retain the outermost protruding points associated with both reference shapes, which will bias the generated shape towards larger areas and perimeters. Since septic real neutrophils exhibit pronounced single cell heterogeneity, the inclusion of a few unusually large cells with this pairwise sampling skewed the shape distribution of septic generated neutrophils towards larger areas and perimeters. In comparison, healthy real and generated neutrophils exhibited no statistical difference in any of the measured shape features, likely since they were more homogeneous in shape. It should be noted that the septic neutrophils could include some subsets that are more dysregulated (perhaps prematurely released from the bone marrow) and others that are phenotypically more similar to healthy neutrophils. If so, the presence of this latter subset could obfuscates the separation of healthy and septic neutrophils by morphology.

### Comparison of real and generated shapes based on primate mandibular molars

As a final case study with three-dimensional shapes, we applied the *α*-shape sampler to a dataset consisting of *N* =15 computed tomography (CT) scans of mandibular molars from two suborders of primates: 8 of these real teeth came from the genus *Microcebus* of the Strepshirine suborder and the remaining 7 came from the *Tarsius* of the Haplorhini suborder ^2,65,66^. The *α*-shape sampler was used to synthetically generate an additional *N ^∗^* = 10 teeth from each genus using the parameters *J* = 2, *θ* = 0, and *ε* = 0.001. In this analysis, we had to set *θ* = 0 because the CT scans for each molar came in the form of boundary meshes, which are technically a “hollowed” representation of fully dense 3D objects (see Materials and methods). This effectively meant that each reference tooth had volumes equal to 0. As a result, we had to avoid setting *θ >* 0 to keep the acceptance probability of new candidate points from being nearly 0 (i.e., we would reject nearly 100% of new candidate points).

It is worth briefly noting that the original dataset started with *N* = 10 *Microcebus* teeth and *N* = 18 *Tarsius* teeth, respectively. Some of these references were removed from the analysis after we estimated their reach values (see, again, the second step in Fig 2b) and observed some distinct outliers which would affect our ability to generate new and realistic shapes downstream. For the *Microcebus* genus, the teeth we used in our analysis had estimated reach values in the range 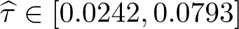, while the unused teeth had values 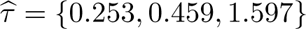 (somewhere 10*×* to 100*×* larger than the rest of the data). Similarly, for the *Tarsius* genus, data for our analysis was restricted to teeth with estimated reach values which fell in the range of 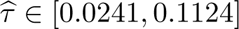, while the omitted teeth had reaches between 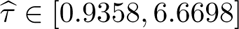. When using all teeth, even with proper alignment and scaling, we generated unrealistic shapes (e.g., synthetic teeth with six or eight roots, which do not occur in either species). A key feature of the *α*-shape sampler is that it allows users to use the estimated 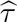 to identify reference shapes that are outliers relative to the rest of input dataset. This can be used to proactively prune reference shapes or use the 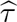 values post hoc to diagnose why the algorithm produced a shape that does not fit with the original set.

A comparison of the quality controlled real teeth and the generated teeth from the *α*-shape sampler can be found in Fig 5a-5d. Overall, we chose this specific collection of molars for our analysis because of the phylogentic relationship between the *Microcebus* and the *Tarsius* (Fig 5e) ^67^. Morphologists and evolutionary anthropologists have previously used this data to understand variations of the paraconid, the cusp of a primitive lower molar. The paraconids do not appear in other genera ^68,69^ and are only retained by *Tarsius* which allows this genus of primate to eat a wider range of foods ^70^. When using these teeth as reference data in our shape generation pipeline, we see that the *α*-shape sampler is indeed able to produce newly generated teeth that qualitatively preserve key features shared between both species (e.g., the four roots) as well as recapitulate species-specific variation that is driven by the presence of the paraconids in the *Tarsius*. More specifically, the generated *Microcebus* teeth are missing the distinguished paraconid that is captured in the generated *Tarsius* teeth (again see Fig 5a-5d), repeating the patterns we see in the real data.

**Figure 5.**
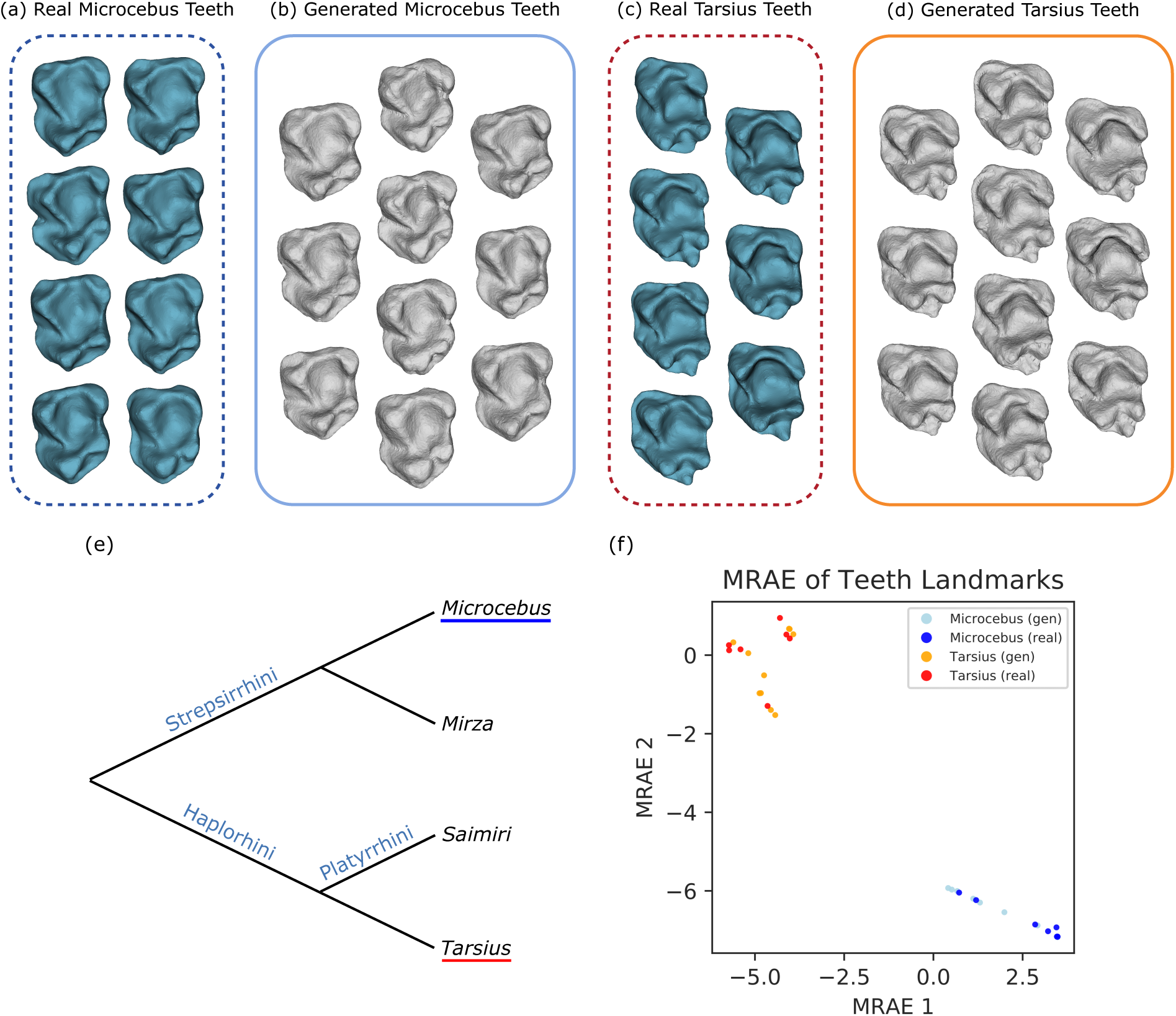
Application of the *α*-shape sampler to generate synthetic 3D primate mandibular molars. Here, we qualitatively compare meshes of **(a)** real *Microcebus*, **(b)** generated *Microcebus*, **(c)** real *Tarsius*, and **(d)** generated *Tarsius* teeth. Morphologically, we know that tarsier teeth have an additional high cusp (highlighted in red) which allows this genus of primate to eat a wider range of foods ^70^. Here, we see that the generated *Tarsius* teeth from the *α*-shape sampler preserve the unique paraconids. In panel **(e)**, we show the phylogenetic relationship between the *Microcebus* and *Tarsius* genus. It has been estimated that the divergence dates of the *Microcebus* and *Mirza* from *Tarsius* happened around five million years before the branching of *Tarsius* from *Saimiri* ^67^. **(f)** We use a manifold regularized autoencoder (MRAE) to show that the generated teeth cluster and intermix with the real *Microcebus* and *Tarsius* teeth, respectively. Figure S4 shows that the same results hold regardless of the dimensionality reduction technique that is used.

To further assess the quality of the shapes produced by the *α*-shape sampler, we follow Turner et al. ^12^ and used Procrustes analysis ^71,72^ to assign 400 landmarks onto each reference and newly generated tooth (Materials and methods). The (400*×*3)-dimensional matrix of landmark points for each shape was reshaped to a scalar vector of length 1200. This was then projected onto a two-dimensional space using the manifold regularized autoencoder (MRAE) on PHATE coordinates (Fig 5f). As expected, we see the real *Microcebus* and the real *Tarsius* teeth form distinctly separate groups along both MRAE1 and MRAE2. We also see the generated *Microcebus* teeth group together with the real *Microcebus* teeth, while the generated *Tarsius* teeth group together with the real Tarsius teeth. These general trends were again confirmed to be independent of the choice of dimension reduction method (Fig S4). For a more quantitative analysis, we also computed the average pairwise Euclidean distance between each tooth group (e.g., Table S4). Here, we observe that the generated *Microcebus* and generated *Tarsius* teeth are nearly twice as close to their respective real groups than to any other group. We attribute the nonzero distance between the generated and real teeth to the fact that we end up accepting all randomly sampled points during our shape generation algorithm (see Materials and methods).

## Discussion

In this paper, we introduced the *α*-shape sampler: a probability-based generative model for two-dimensional and three-dimensional shapes. The underlying theoretical innovation of connecting the mathematical concept “reach” with the *α* parameter in *α*-shapes allows us to implement a data-driven algorithm with the scalability to accommodate the growing sizes of emerging imaging and shape-based databases. We applied our generative pipeline to both 2D and 3D datasets and demonstrated its ability to successfully capture important geometric, morphometric, and topological characteristics of complex objects. In the main text, we focus on demonstrating our generative model when reference shapes are available. This is meant to approximate the reality that the underlying manifold for shapes observed in many biological applications is often unknown. In the Supporting Information, we derive theory and discuss how to generate new shapes when the true manifold is indeed known and available (Fig S5-S12). This includes detailing how one might sample new shapes directly from probability distributions (code for this “exact” approach is also included in our open-source R package; see URLs).

The current implementation of the *α*-shape sampler framework offers many directions for future development. For example, there are a few considerations to be made when choosing the *J* number of reference shapes and the *θ* threshold for accepting new candidate points in the *α*-shape sampler pipeline. Almost counter-intuitively, the smaller we select *J* to be, the more variation there will be in the generated shapes. This is because the joint point cloud starts to converge as the number of *J* shapes that are included grows. Additionally, the number of *J* reference shapes limits the number of new shapes that can be produced. Combinatorially, we can only generate 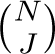 new shapes. While this may be seen as a limitation, it also prevents us from augmenting a study with generated shapes that are too far outside of what has been observed in real data. Similarly, when selecting *θ*, our suggestion is to choose *θ* = *d* (the dimension of the shape space) so that one avoids noisy points and edges around the boundary. The exception to this rule is when the reference shapes are in the form of boundary meshes which are technically a lower dimensional representation of the full shape data. For example, the primate teeth meshes analyzed in the main text are two-dimensional simplices in three dimensions. In this case, we recommend *θ* = 0 such that all points are accepted. While this removes the possibility for noise and variation between iterative runs of the *α*-shape sampler, even choosing *θ* = 1 will result in such a strict threshold of acceptance that the new shape will be a few isolated points scattered in space. We believe this happens because the volume of intersection of a two-dimensional surface mesh with a three-dimensional ball is 0 due to the mesh having Lebesgue measure 0. While the generated shapes may end up being thicker meshes, this can be fixed via post-processing of the data. To avoid this issue, it is best to use shapes that are “filled” in (such as the neutrophil example), but sometimes this is not feasible or practical for the given dataset.

In its current form, the *α*-shape sampler performs considerably better when the reference shapes in the input dataset are well aligned. Indeed, alignment was performed with the simulated annuli and tori (Fig 3), as well as with the mandibular molars which included landmarks amenable to unsupervised learning methods (Fig 5). In comparison, neutrophil morphologies lacked such landmarks and so shapes were only centered on their centroids (Materials and methods). Nevertheless, real and generated shapes for healthy neutrophils were statistically similar, since the real morphologies exhibited comparable areas and were relatively compact (Fig 4). However, some generated shapes for septic neutrophils considerably exceeded the corresponding real shapes in area and perimeter, since the *α*-shape sampler generates manifolds that retains the outermost protruding points associated with both shapes (Table S3). To address this artifact, we attempted to rescale shapes after generation to match areas and perimeters, which distorted circularity and convexity. Alternatively, aligning neutrophils along their long axis tended to bias towards the generation of more elongated morphologies. It is conceivable that septic neutrophils with very different morphologies belong to different subsets, and so the generated cell is a chimera based on different subsets without a plausible biological basis. These issues could be addressed in highly heterogeneous populations by sampling a larger number of single cells to limit the biasing effect of outliers, and to discard any generated cells that deviate excessively from the real shape distribution. Future work could also utilize additional information based on cell migration or tractions ^60,73,74^, along with single-cell genomics ^75^ to gain additional insight into septic cell phenotype. Finally, this approach could be effective for other cell types, such as analyzing the epithelial-mesenchymal transition, since the associated spindle-like morphology displays more consistent landmarks for shape alignment^76–78^.

From a statistical perspective, the assumption that all points in the input data point clouds are uniformly distributed over the same underlying manifold may not be suitable for all applications. When points are not uniformly distributed, the calculation of reach becomes less precise because there is too much variance between boundary points. As a result, the *τ* estimate ends up too big in some parts of the point cloud and too small in others, leading to the loss of local geometric information and the possible addition of global topological information, both of which hinder the ability to generate new realistic shapes that properly fit in the same class as the input dataset. Where points are not uniformly distributed, it may be the case that *α*-shapes are the appropriate tool for modeling shapes, as was studied in Gerritsen ^79^. This is particularly true when points have additional contextual meaning (e.g., molecular structures such as proteins or strands of DNA) or in cases where meshes are very detailed in some areas and less so in others. An immediate future avenue of work is to extend our pipeline to work for weighted *α*-shapes ^80^, coupled *α*-shapes ^81^, and *β*-shapes ^82^ to fit a broader range of applications.

### URLs

Code for the *α*-shape sampler and data simulations is available at https://www.github.com/lcrawlab/ ashapesampler. Slicer auto3dgm paradigm is available at https://toothandclaw.github.io/. Binary masks of the healthy and septic neutrophils and 3D meshes of the primate mandibular molars are available on the Harvard Dataverse at https://doi.org/10.7910/DVN/K9A0EG. Scripts to reproduce the results in this paper are also publicly available and can be found at https://github.com/lcrawlab/ashapesampler_paper_results.

## Materials and methods

### Introduction on *α*-shapes

In this work, we consider a shape to be the simplicial complex approximation of a compact Riemannian manifold embedded in Euclidean space. We use the same definitions for simplices and simplicial complexes as presented in Edelsbrunner and Harer ^83^. We also assume that all shapes considered in a given phenotypic class (e.g., healthy septic cells or molars from a given species of primate) have vertices sampled from the same underlying manifold and that the variation observed across shapes within the class stems from a finite sampling of points. When we know the true underlying manifold, we can generate shapes using hierarchical probability distributions (see Supporting Information). The demonstration of the *α*-shape sampler in the main text (and what we detail throughout this section) demonstrates how we can generate new shapes when we have data instead of the underlying manifold. Given our applications in the main text, we will derive the details of our probabilistic generative framework while assuming that we are working with shapes that are *d* = 2 or 3 dimensions; however, also note that the theory we present is generally applicable to larger finite dimensions as well.

We define *α*-shapes using Voronoi cells and the Deluanay triangulation. The main motivation behind this choice is that it mirrors how we compute *α*-shapes in practice and we believe that this construction provides a more intuitive framing for understanding the parameters in our sampling algorithm. For a more rigorous definition, we refer the reader to Edelsbrunner et al. ^38^. To begin, we assume that all points are in general position. That is, in the *d*-th dimension ^84^, we assume the following:

- No *d* + 1 points are colinear or coplanar;
- No *d* + 2 points are cocircular or cospherical;
- No points form a smallest circle or cicumsphere of radius *α*;
- No points lie on the smallest circumsphere of *d* + 1 other points.

In practice, this assumption is relatively strict and rarely occurs naturally; however, in the Supporting Information, we prove that this assumption holds true in our generative algorithm so long as points are sampled uniformly. In real data applications, users can either ignore the points during the estimation of reach *τ* (e.g., as we do with the primate mandibular molars) or perturb the points slightly to correct for this assumption (e.g., as we do with the segmented images of the neutrophils).

Let *S* denote a set of *P* points in ℝ*^d^* in general position. The *Voronoi cell* of a point *p ∈ S* is the set of points in ℝ*^d^* for which *p* is the closest. We denote the Voronoi cell as the following

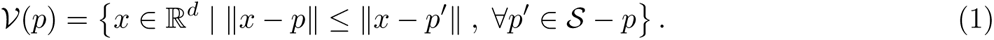

The *Voronoi diagram* of *S* is then the union of all Voronoi cells and takes up the space of ℝ*^d^*. The *Delaunay complex* of *S* is isomorphic to the nerve of the Voronoi diagram. As long as the points of *S* are in general position, the Delauany complex of *S* is well-defined and forms the convex hull of the points *S* in R*^d^*. This is often referred to as the *Delaunay triangulation* of *S* and is denoted by

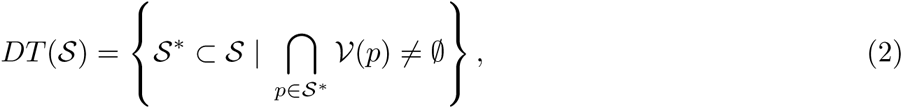

where *S^∗^*is a subset of points in *S* and ∅ represents the empty set. The example in Fig 1 depicts the Delaunay triangulation and the convex hull for a point set. Instead of Voronoi cells which together take up the entire space, we can look at subsets of those cells. Let *B_α_*(*p*) denote a ball of radius *α* centered at point *p*. Furthermore, let *R_p_*(*α*) = *B_α_*(*p*) *∩ V*(*p*) denote the intersection of the Voronoi cell of *p* and the ball of radius *α* centered at *p* (e.g., see the gray shapes in Fig 1). The union of *R_p_*(*α*) for all points *p ∈ S* form a cover of *S*, the nerve of which forms the *α-complex* which we will denote as *S_α_*. The boundary of *S_α_* defines the *α-shape*. Formally, the border is defined by *α-extreme* points, which are the points *p^∗^ ∈ S* such that there exists a ball of radius *α* with *p^∗^* on the border where the complement of the disc contains all other points in *S*. In Fig 1a-c, we see that all points are *α*-extreme; while in Fig 1d, we see that *α* becomes large enough such that one point is not *α*-extreme and is therefore an *interior* point of the shape. Finally, in Fig 1e, there are three interior points and the rest are boundary or *α*-extreme points.

### Estimating the reach parameter *τ*

Assume that we have a dataset with *N* shapes or images. We will refer to these samples as “reference shapes” from which we will generate new shapes. Let *K_i_* = *{V_i_, E_i_, F_i_, T_i_}* denote the mesh for the *i*-th observation in the reference set comprised of a collection of vertices *V_i_*, edges *E_i_*, faces *F_i_*, and tetrahedra *T_i_* (if applicable). Recall that (i) we assume that all vertices for reference shapes in the same phenotypic class come from the same underlying manifold, and (ii) most real shape and imaging data do not readily come in the form of *α*-shapes or *α*-complexes. In order to generate new shapes, we must derive an appropriate point set from the reference shapes (both in terms of location in space and in the total number of vertices) and we must find an appropriate value of *α*. To do so, we use the concept of *reach* (denoted by *τ*) as presented in Aamari et al. ^39^, which can also be related to the inverse of the condition number as introduced in Niyogi et al. ^85^ (see Supporting Information for a formal definition). In practice, *τ* is the minimum distance from the boundary of a shape to its medial axis and can be approximated as either the minimum distance between connected components or the minimum radius of any holes (or voids) in a shape.

At a high level, we estimate the reach *τ_i_* for the *i*-th reference shape by using the boundary points of its simplicial complex *p ∈ ∂K_i_* (i.e., the *α*-extreme points in an *α*-shape). We do this because the boundary information is all that is relevant to estimating reach. The collection of ***τ*** = (*τ*_1_*, …, τ_N_*) values from the *N* reference shapes are then used to estimate an appropriate value of *α* for the newly generated shapes. Other theoretical methods for estimating reach using an underlying manifold have been proposed ^39,86,87^, but we use this approximate estimate to optimize computational speed. By connecting *α* to *τ*, we ensure the preservation of major topological and geometric characteristics for the simplicial complex derived from the *α* parameter over a point set. The reach estimates *τ* can also be used to sample a point set for the new shapes, both in point set size (i.e., how many vertices we need to sample from the underlying manifold) and in point density. We substitute the minimum number of points needed to preserve the homology of the underlying manifold with an *α*-dense cover using the main result in Niyogi et al. ^85^ (Supporting Information). Once *τ* is derived from the input reference dataset, the appropriate *α* can be selected and a new point set can be sampled—the combination of which will allow use to generate new shapes.

Algorithmically, the process of estimating the reach *τ_i_* for the *i*-th reference shape is done using the following procedure.

- Examining a boundary vertex *p ∈ ∂K_i_*, we first learn its distance to neighboring sets of vertices *q ∈ N_i_*(*p*) by studying the corresponding edges *E_i_* that are present in the mesh. We save the largest of these distances using the Euclidean distance, *d_E_* = max*_q∈Ni_*_(_*_p_*_)_ *p − q*.
- Next, we define *C_p_* to be the set of circumcenters of all faces in *F_i_* and tetrahedra *T_i_* containing *p*. These circumcenters are the points at which any three or four points would meet in the Voronoi diagram and, hence, where faces and tetrahedra would form in the resulting *α*-complex. We also save the largest of these distances *d_C_* = 2 max*_c∈Cp_ p − c*. Here, we take twice the value of the circumcenter distance in an effort to preserve consistency across dimensions. Recall that for *d_E_*, we consider the entire lengths of edges, not just the midpoints. The circumcenter can be interpreted as a rough estimate of a “midpoint” for faces and tetrahedra; as a result, we multiply that value by 2 to capture the full “distance” *d_C_*.
- Once we have these two distances corresponding to edges and circumcenters involving point *p*, we take the maximum which we denote as *d_p_* = max(*d_E_, d_C_*). Each value *d_p_* indicates how large *α* needs to be in order to recover the geometric properties in a localized region of the reference mesh.
- In practice, we find the next furthest point outside of the minimum *d_p_* range because it establishes the largest that *α* can be without us losing any geometric information. To do so, we consider the set of vertices in *V_i_* that do not share an edge with *p* but are more than *d_p_* distance away. Formally, this set is 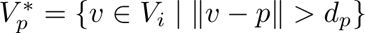. The *τ* value for a given point is computed as

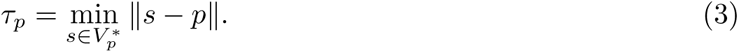

In the event that 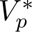 is empty (e.g., when *p* shares an edge, face, or tetrahedra with all other vertices in *V_i_*), we take *τ_p_* = *d_p_*.

- The reach for the *i*-th mesh shape is approximated by

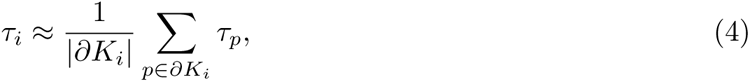

which is the mean *τ* value for all boundary points in the shape where *|∂K_i_|* denotes the cardinality of the set.

Note that other summary statistics could be used in the final step, such as taking the minimum *τ_p_* across all points, but empirically we find that taking the mean gives robust estimates and keeps outliers from artificially deflating the value of *τ_i_*. For example, in theory, the true reach estimate would take the minimum of *τ_p_* over all boundary points; however, a small outlier *τ_p_* value would lead to a small *τ_i_* when we take the minimum and that would result in computational bottlenecks when we later generate shapes.

Therefore, we choose to trade the precise theoretical implementation for computational scale without compromising major shape information. Repeating this procedure for all *N* meshes in the dataset yields a collection of estimated reach parameters ***τ*** = (*τ*_1_*, …, τ_N_*) which we will use to generate new point clouds and shapes.

### Algorithm for generating new shapes

When generating new shapes, the first task is to create a corresponding point cloud. This step requires developing a method for sampling points from some underlying manifold *M*. Ideally, one could fit a function to each reference shape from a given dataset, average the functions to approximate the true manifold, and then sample new points directly from that manifold via rejection sampling to simulate uniformity. This strategy is similar to what Diaconis et al. ^21^ illustrates on the torus; however, this same approach is computationally infeasible for modern datasets with tens to hundreds of shapes. One could use techniques from manifold learning to generate point clouds, but the available techniques involve black-box methods such as dual generators ^25^ and autoencoders ^27^. While these approaches have been shown to be useful for assessing predictive models, these do not provide enough interpretability to learn much about the underlying functional representation of the manifold. We could recover a function for each shape using Gaussian processes, as what is done in Albrecht et al. ^13^, but to practically implement this strategy, we need to have access to landmarks for each shape. Once we have our point set, we need to find an *α* parameter for the shape to dictate how to reconstruct the shape. Most imaging and shape datasets will not be in the form of *α*-complexes as the points in many applications are not in general position. As a result, we need an algorithm that can give us both an accurate point cloud from the underlying sub-manifold and the correct parameter for constructing the *α*-shape.

Sampling uniformly from balls with radius 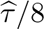 around points in a given reference point cloud allows us stay close to what we assume to be the true manifold without directly calculating the manifold itself. Additionally, while this procedure is not exactly the same as uniform sampling (i.e., points that are closer together will have balls with greater overlap), we conjecture that the overall sampling ends up matching the true density of the point set. Adding a rejection-like step to the sampling scheme then gives the algorithm robustness to outlying points or atypical features that are present in shapes from the reference dataset. We will work with the “approximate manifold” given by the union of balls around the corresponding reference point clouds of radius 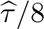; call this manifold 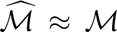. In practice, we avoid estimating or calculating the underlying manifold, but we stay true to the given reference data by implementing a “rejection sampling-like” algorithm via the following five step procedure:

1. Choose 2 *≤ J ≤ N* number of reference shapes from the input dataset to serve as references and combine their corresponding point clouds into a joint set denoted as *Q*.
2. Determine the number of candidate points **y** to sample based on a ball of radius 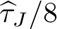 centered around reference points 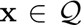. Here, 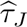 is the minimum value in ***τ*** corresponding to the subset of *J* selected reference shapes. Note that this 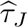 value will change depending on the subset of *J* reference shapes chosen for the generation of new shapes. The variation of 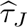 across different subsets of reference shapes contributes to the variation observed in newly generated shapes.
3. Given a reference point *x ∈ Q* in the joint point cloud of the *J* reference shapes, sample random candidate points **y** from 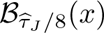— that is, sample random points **y** from a small ball of radius 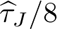 centered at point *x*.
4. Calculate the number of additional points in the joint point cloud *z ∈ Q* that lie within a ball centered at each candidate point *y* which we define as 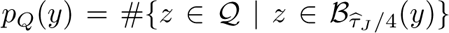. This number does not include the original reference point *x* from the previous step. Next, choose *θ ≤ p_Q_*(*y*) to be the minimum number of points needed to accept each new candidate point *y*. This sets up the following accept-reject decision rules for the generation of new shapes where:

- If *p_Q_*(*y*) *≥ Jθ*, accept point *y*.
- If *p_Q_*(*y*) *< Jθ*, accept point *y* with rate 1 *−* exp*{−*2(*p_Q_*(*y*) *− θ*)*/Jθ}*.
- If *p_Q_*(*y*) *< θ*, reject point *y*.

We detail the logic behind this rejection rule below.
5. Repeat these steps for all points in the combined point cloud *x ∈ Q*.

There are a few key takeaways in the procedure specified above. First, we sample new points uniformly from one ball at a time rather than from the union of balls. This means that the new point cloud will reflect the density of the combined point cloud *Q* from the subset of *J* reference shapes. Second, to add some variance to the sampled point cloud and to ensure confidence in the newly sampled points, we implement the following rejection-like rule:

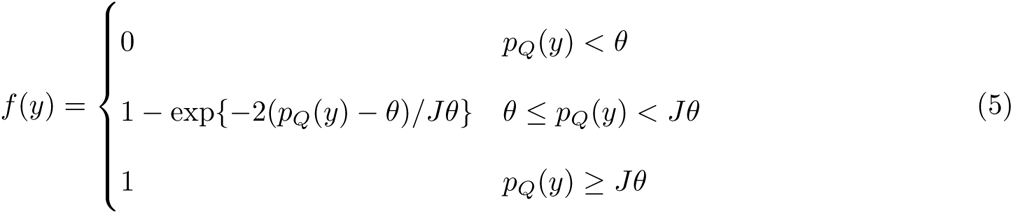

where, again, *p_Q_*(*y*) is the number of points in the joint point cloud *Q* that are within a 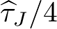 radius of the candidate point *y*; *θ* is the minimum number of points we require from the reference point cloud *Q− x* to be within a 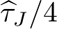 radius of the candidate point *y* in order to accept *y* (as the reference point *x* is already within that radius by definition); and *J* is again the number of reference shapes. Note that in Eq (5), the choice of *J* will affect the rate of acceptance and will approach 1 as *p_Q_*(*y*) *→ Jθ*. The three-part rule in Eq (5) is designed to accommodate three scenarios when we consider to accept a newly sampled point *y*. If *p_Q_*(*y*) *< θ*, then there are fewer neighboring reference points than desired and indicates that the candidate point *y* is likely to be far away from the boundary of the point cloud. We have little confidence that these points are from the manifold that we wish to approximate 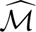 and so, consequently, we reject these points. In the scenario where *p_Q_*(*y*) *≥ Jθ*, the candidate point *y* is near more than *θ* real points (on average) from the *J* reference shapes. In this case, we have high confidence that *y* is from the approximated manifold 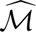 and automatically accept it as a newly sampled point.

In the middle scenario, where *θ ≤ p_Q_*(*y*) *< Jθ*, we want a rule that allows for some uncertainty in *y* as a function of the number of nearby points *p_Q_*(*y*) from the *J* reference shapes. Here, we choose 1 *−* exp*{−*2(*p_Q_*(*y*) *− θ*)*/Jθ}*, which is the cumulative distribution function (CDF) for an exponential random variable with rate *Jθ/*2 that is shifted to be 0 when *p_Q_*(*y*) = *θ* (i.e., the threshold for the minimum number of points needed to accept each new candidate point *y*). The exponential distribution is typically used to model the amount of time until some specific event occurs—where there are fewer large values and more small values. The main motivation behind this choice is to reward candidate points *y* that with higher values of *p_Q_*(*y*). When we have *J* = 2 reference shapes, the rate of the distribution will be *θ*; as we add more reference shapes to the algorithm, the rate at which we find more neighboring points for any candidate point *y* will increase. In practice, using our proposed rejection-like rule, the acceptance rate will be roughly 100% for randomly drawn candidate points that are near the interior of the point cloud (particularly in regions where the *J* reference shapes being used all overlap). Intuitively, the rate of acceptance will decrease for new candidate points that are sampled near the boundaries of the *J* reference shapes. The range of the overall acceptance probability will depend on the intraclass heterogeneity of the reference dataset and the quality of alignment of the point clouds during preprocessing.

### Patient blood sample collection and primary neutrophil isolation

Blood was drawn from healthy donors or septic patients with written informed consent at Rhode Island Hospital, in accordance with the guidelines and approval of the Institutional Review Board. Briefly, healthy donors had no known acute infection or chronic systemic disease within one month prior to the blood draw. We did not collect blood from minors, pregnant women, prisoners, mentally retarded or mentally disabled patients or volunteers. Septic patients from the surgical intensive care unit (ICU) and the trauma ICU displayed at least two systemic inflammatory response syndrome criteria with a source of infection, and enrolled within 48 hours of their diagnosis or admission. Patients also had to be at least 18 years of age without a massive blood transfusion. Further details on study design are documented elsewhere ^60^.

For both healthy donors and septic patients, 10-30 milliliters (mL) of blood was collected in EDTA-containing Vacutainer tubes. Buffy coat was separated by centrfiguation with Histopaque-1077 with an additional sedimenation step for neutrophils using 3% Dextran (400-500 kDa). Any contaminating erthrocytes were eliminated by hypotonic lysis, and neutrophils were then resuspended in cation-free HBSS media.

### Polyacrylamide gel preparation and neutrophil imaging

Briefly, polyacrylamide gel substrates were polymerized on a 25 millimeters (mm) glass coverslip, using 3% acrylamide and 0.2% bisacrylamide for a Young’s modulus of E = 1.5 kPa, along with fluorescent red 0.5 *µ*m carboxylate-modified polystyrene beads. Gel substrates were then coated with human fibronectin (Gibco 33016015) using the photoactivatable crosslinker sulfo-SANPAH (Sigma 803332) and rinsed extensively. Further experimental details are documented elsewhere in Oakes et al. ^73^ and Witt et al. ^60^, respectively. The polyacrylamide gel and coverslip were mounted in a coverslip holder, then covered with 1 mL of Leibovitz L-15 media. About 50,000 neutrophils were plated and allowed to adhere for 15 minutes. Approximately 20-60 adherent cells were imaged in phase microscopy using a Nikon TI-2 epifluorescent microscope using a 40X air objective with a 0.6 numerical aperture. An Okolab enclosure around the TI-2 maintained the apparatus at 37*^◦^* and 5% CO_2_ for the duration of the experiments. Only adherent cells were selected for imaging. The *N* represents the number of individual neutrophils imaged and analyzed, with an *n >* 3 for individual septic or healthy donors.

### Converting segmented neutrophil images to 2D simplicial complexes

To convert tif files into two-dimensional simplicial complexes, we used a multi-step procedure. For the healthy neutrophils, each image was first cropped to include only the middle 50%. Septic neutrophil images were already cropped. Next, the centroid of each shape was found using the median row and column; cells were centered by placing this centroid at the center of the new matrix. The black-and-white cell images were converted into a binary matrix representing black-and-white pixels. This matrix was then searched to find all the black pixels, which were used as vertices for the complex. To add randomness to the pixel points, all vertices were also perturbed within their pixel areas. Next, edges were formed by finding pairs of vertices that were either orthogonally or diagonally adjacent according to the matrix. However, in order to avoid overlapping edges, the upper left and downward right diagonals of each vertex were removed except when upper right and downward left diagonals could not exist (such that the overlap would be impossible). Finally, every three edges that could form a triangle were listed as a face to construct a group of adjacent faces, which was plotted to generate a 2D simplicial complex for the image.

### Evaluation of generated neutrophils

Representative cell morphologies were manually traced, converted to binary masks, and then turned into simplicial complexes (Fig 2a). The *α*-shape sampler was used to synthetically generate additional cells with parameters *J* = 2 and *θ* = 2. These newly generated neutrophils were then converted to binary masks. We computed 33 geometric characteristics using the masks of the original and the generated shapes, respectively, including: area, perimeter length, number of protrusions, compactness, and others as described in Bhaskar et al. ^61^. The vectors of these characteristics were projected onto a two-dimensional latent space using a manifold regularized autoencoder (MRAE) ^62^ where the loss function is the combination of a mean square error loss on the autoencoder itself and the “Potential of Heat-diffusion for Affinity-based Transition Embedding” (PHATE) coordinates in latent space. This combined loss function is formally defined as the following

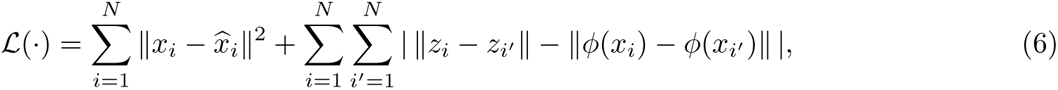

where *L*(*·*) denotes the loss function associated with the autoencoder; *N* is the number of shapes in the dataset; *x_i_* is the input data for the *i*-th shape; ‖*·*‖ is the *L*^2^-norm; 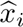 is the reconstructed version of the *i*-th shape as determined by the decoder portion of the MRAE; *z_i_* is the two-dimensional latent embedding for the the data associated with the *i*-th shape; and *φ*(*·*) is the PHATE function mapping the shape data to ℝ^2^. The idea behind the loss function is to train an autoencoder to not only minimize the difference between the input and reconstructed data, but also force the latent space to behave as similarly as possible to the PHATE function *ϕ*. Since PHATE is a dimensionality reduction method designed to honor the original local and global structure of high-dimensional data ^64^, adding the extra loss component based on the PHATE coordinates in the latent space forces the autoencoder to also honor the original structure of the data as well.

In addition to the MRAE, we also assess the new shapes generated by the *α*-shape sampler using other dimensionality reduction approaches including: the uniform manifold approximation projection (UMAP) ^63^, PHATE, principal component analysis (PCA), and a generic autoencoder. Each of these analyses were used to demonstrate that our conclusions about the shapes produced by the *α*-shape sampler are robust regardless of the unsupervised dimension reduction method that we choose. Briefly, UMAP was implemented with 5 nearest neighbors, 2 connected components, Euclidean distance, and a minimum distance set to 0.1. PHATE was implemented with 5 nearest neighbors, 2 connected components, a Von Neumann Entropy diffusion operator, log potential, Euclidean distance, and we used stochastic gradient descent for the multi-dimensional scaling method. Both the autoencoder and the MRAE were trained with 500 epochs.

### Evaluation of generated primate manibular molars

To generate synthetic primate manibular molars, we used parameters *J* = 2 and *θ* = 0 in the *α*-shape sampler software, which meant an automatic 100% acceptance rate of sampled points. Since the reference teeth data were given as two-dimensional surface meshes in three-dimensional space, they had volumes equal to 0. In this case, setting *θ >* 0 would send the acceptance probability of new candidate points to nearly 0 (i.e., we would reject nearly 100% of new candidate points). Our evaluation for the generated shapes with this dataset were similar to the landmarking and subsequent dimensionality reduction analyses used in Turner et al. ^12^. First, the reference teeth were aligned using the sofware pacakge auto3dgm ^88^. We then generated 10 new synthetic teeth each from the *Microcebus* and the *Tarisus* genera, respectively. We used Procrustes analysis ^71,72^ to assign 400 landmarks to each newly generated tooth so that these could also be aligned and scaled. The (400*×*3)-dimensional matrices of landmark points for both the newly generated and real reference teeth were reshaped to scalar vectors of length 1200. These were then projected onto a two-dimensional space using the same manifold regularized autoencoder (MRAE) and other dimensionality reduction techniques (UMAP, PHATE, PCA, and an autoencoder) as was done the neutrophils. UMAP was implemented with 5 nearest neighbors, 2 connected components, Euclidean distance, and a minimum distance set to 0.1. PHATE was implemented with 5 nearest neighbors, 2 connected components, a Von Neumann Entropy diffusion operator, log potential, Euclidean distance, and we used stochastic gradient descent for the multi-dimensional scaling method. Both the autoencoder and the MRAE were trained with 500 epochs. For quantitative results, we calculate Euclidean distances between the length 1200 scalar vectors representing each tooth and gather the pairwise distances to reaffirm that the generated teeth are appropriately spaced from the original reference datasets.

## Acknowledgements

This research was conducted using computational resources and services at the Center for Computation and Visualization (CCV), Brown University.

## Author Contributions

- **Conceptualization:** Emily T. Winn-Nuñez, Ian Y. Wong, Lorin Crawford
- **Data Curation:** Hadley Witt, Jonathan S. Reichner, Lorin Crawford
- **Formal Analysis:** Emily T. Winn-Nuñez
- **Funding Acquisition:** Emily T. Winn-Nuñez, Hadley Witt, Dhananjay Bhaskar, Jonathan S. Reichner, Ian Y. Wong, Lorin Crawford
- **Investigation:** Emily T. Winn-Nuñez, Lorin Crawford
- **Methodology:** Emily T. Winn-Nuñez, Dhananjay Bhaskar, Lorin Crawford
- **Project Administration:** Lorin Crawford
- **Resources:** Jonathan S. Reichner, Ian Y. Wong, Lorin Crawford
- **Software:** Emily T. Winn-Nuñez
- **Supervision:** Jonathan S. Reichner, Ian Y. Wong, Lorin Crawford
- **Validation:** Emily T. Winn-Nuñez, Ryan Y. Huang, Jonathan S. Reichner, Ian Y. Wong, Lorin Crawford
- **Visualization:** Emily T. Winn-Nuñez, Ryan Y. Huang, Ian Y. Wong, Lorin Crawford
- **Writing - Original Draft:** Emily T. Winn-Nuñez, Ian Y. Wong, Lorin Crawford
- **Writing - Final Draft:** Emily T. Winn-Nuñez, Hadley Witt, Dhananjay Bhaskar, Ryan Y. Huang, Jonathan S. Reichner, Ian Y. Wong, Lorin Crawford

