## Supplementary Text and Figures for "Generative modeling of biological shapes and images using a probabilistic *α*-shape sampler"

### Supporting Information to "Generative modeling of biological shapes and images using a probabilistic $\alpha$-shape sampler"

Emily T. Winn-Nuñez[1,†], Hadley Witt[2,3], Dhananjay Bhaskar[4], Ryan Y. Huang[5], Jonathan S. Reichner[2,3], Ian Y. Wong[6], and Lorin Crawford[7-9†]

**1 Division of Applied Mathematics, Brown University, Providence, RI, USA**

**2 Graduate Program in Pathobiology, Brown University, Providence, RI, USA**

**3 Division of Surgical Research, Department of Surgery, Rhode Island Hospital, Providence, RI, USA**

**4 Department of Genetics, Yale School of Medicine, New Haven, CT USA**

**5 Department of Computer Science, Brown University, Providence, RI USA**

**6 School of Engineering, Legoretta Cancer Center. Brown University, Providence, RI USA**

**7 Microsoft Research, Cambridge, MA, USA**

**8 Department of Biostatistics, Brown University, Providence, RI, USA**

**9 Center for Computational Molecular Biology, Brown University, Providence, RI, USA**

**† Corresponding**

### 1 Background and Theory

In this section, we give a thorough theoretical background on $\alpha$-shapes in the topological sense and discuss how they can be considered in applications related to generative modeling in biology. We discuss the merits of using alpha-shapes compared to other shape reconstruction methods and lay the theoretical foundations for our $\alpha$-shape sampling algorithm when we have access to the true underlying manifold. Throughout the main text and this supplementary text, when we refer to a *shape*, we mean a simplicial complex where the points are sampled from a compact sub-manifold embedded in $d$-dimensional Euclidean space. Note that we focus on applications where $d = 2$ or 3 in our study, and we leave theoretical extensions to higher dimensions and/or non-Euclidean spaces for future work.

#### 1.1 Formal definition of $\alpha$-shapes

We use the same definitions for simplices and simplicial complexes as in Edelsbrunner and Harer (2010). Following the notation from the main text, let $\mathcal{S}$ denote a set of $P$ points in $\mathbb{R}^d$ (where $d = 2$ or 3) in general position. The *Voronoi cell* of a point $p \in \mathcal{S}$ is the set of points in $\mathbb{R}^d$ for which $p$ is the closest. The Voronoi cell is formally defined as the following

$$\mathcal{V}(p) = \left\{x \in \mathbb{R}^d \mid \|x - p\| \leq \|x - p'\| \text{ , } \forall p' \in \mathcal{S}\right\}.$$

The *Voronoi diagram* of $\mathcal{S}$ is then the union of all Voronoi cells and takes up the space of $\mathbb{R}^d$. The *Delaunay complex* of $S$ is isomorphic to the nerve of the Voronoi diagram. As long as the points of $\mathcal{S}$ are in general position, the Delauany complex of $\mathcal{S}$ is well-defined and forms the convex hull of the points $S$ in $\mathbb{R}^d$. This is often referred to as the *Delaunay triangulation* of $\mathcal{S}$ and is denoted by

$$DT(\mathcal{S}) = \left\{\mathcal{S}^* \subset \mathcal{S} \mid \bigcap_{p \in \mathcal{S}^*} \mathcal{V}(p) \neq \emptyset\right\}, \tag{1}$$

where $\mathcal{S}^*$ is a subset of the point set in $\mathcal{S}$ and $\emptyset$ represents the empty set. Instead of Voronoi cells, which together take up the entire space, we can look at subsets of those cells. Let $\mathcal{B}_\alpha(p)$ denote a ball of radius $\alpha$ centered at point $p$. Furthermore, let $\mathcal{R}_p(\alpha) = \mathcal{B}_\alpha(p) \cap \mathcal{V}(p)$ denote the intersection of the Voronoi cell of $p$ and the ball of radius $\alpha$ centered at $p$. The union of $\mathcal{R}_p(\alpha)$ for all points $p \in \mathcal{S}$ form a cover of $\mathcal{S}$, the nerve of which forms the *$\alpha$-complex* which we will denote as $\mathcal{S}_\alpha$. The boundary of $\mathcal{S}_\alpha$ defines the *$\alpha$-shape*.

Formally, the border is defined by $\alpha$-*extreme* points, which are the points $p^* \in \mathcal{S}$ such that there exists a ball of radius $\alpha$ with $p^*$ on the border where the complement of the disc contains all other points in $\mathcal{S}$. Two $\alpha$-extreme points $p^*$ and $q^*$ are $\alpha$-*neighbors* if there exists a ball of radius $\alpha$ which has $p^*$ and $q^*$ on the border and contains no other points of $\mathcal{S}$. Given a set of points $\mathcal{S}$ and a real $\alpha > 0$, the $\alpha$-*shape* of $\mathcal{S}$ can be equivalently defined as the straight line graph whose vertices are the $\alpha$-extreme points and whose edges connect the respective $\alpha$-neighbors. In Fig 1a-c in the main text, we see that all points are $\alpha$-extreme; while in Fig 1d, we see that $\alpha$ becomes large enough such that one point is not $\alpha$-extreme and is therefore an *interior* point of the shape. Finally, in Fig 1e, there are three interior points and the rest are boundary or $\alpha$-extreme points. An extension of this figure showing different $\alpha$-shapes being formed as a function of the number of points $P$ sampled from a unit square and the parameter $\alpha$ can be found in Fig S1. Note that when $\alpha = \infty$, the resulting $\alpha$-complex is the closest point Delaunay triangulation complex of $S$. In Edelsbrunner et al. (1983), the authors prove that the $\alpha$-complex is a subcomplex of the closest point Delaunay complex.

Before we detail the general framework for generating shapes from a probabilistic model, we need to first ensure that the $\alpha$-shape being sampled is well-defined. In particular, the Delaunay complex (and thus the $\alpha$-complex, a subcomplex of the Delaunay complex) falls apart when there are either colinear/coplanar or cocircular/cospherical points. As illustrated in Fig S5 in two dimensions, Delaunay tringulation algorithms forming the Delaunay complex assign faces by (i) checking all possible combinations of three points within a point set $\mathcal{S}$, (ii) checking the unique circle defined by those points, and (iii) checking whether those circles contain any other points. Three points make a face if their unique circle does not contain any other point in the point set. If three points are colinear, no such circle (sphere) is defined; and if four points are cocircular, then there are multiple faces that could be selected among the set (Fig S5b). The same is true in three dimensions when four points are coplanar or five points are cospherical. While many algorithms correct for collinearity or cocircularity by perturbing points (e.g., Fig S5c and S5d), avoiding this step saves considerable computational time and an extra layer of consideration for our generative pipeline. For our benefit, we want to show that the probability of this occurrence is 0. As stated in Edelsbrunner and Mücke (1990), there are four requirements to guarantee that the points of a set $\mathcal{S}$ are in general position in $\mathbb{R}^d$:

1. No $d + 1$ points are colinear or coplanar;

2. No $d + 2$ points are cocircular or cospherical;

3. No smallest circumsphere for $k$ points, where $2 \leq k \leq d + 1$, has radius equal to exactly $\alpha$;

4. No points lie on the smallest circumsphere of two or three other points.

We will prove that the degenerate scenarios described above almost surely do not occur. The key to the proof relies on the fact that the Lebesgue measure in $\mathbb{R}^d$ is 0 on subspaces with dimension less than $d$, and that the Lebesgue measure is translation invariant(Rudin, 2013). Therefore, any lines and transformations of $\mathcal{S}^1$ have measure 0 in $\mathbb{R}^2$ and $\mathbb{R}^3$, and any planes and transformations of $\mathcal{S}^2$ have measure 0 in $\mathbb{R}^3$. The uniform measure on a compact Riemannian manifold $\mathcal{M}$ embedded in $\mathbb{R}^d$ is derived from the Lebesgue measure $\lambda$ where, for a subset $\mathcal{A} \subset \mathcal{M}$, the probability a point $x$ lies in $\mathcal{A}$ is given by:

$$\mathbb{P}(x \in \mathcal{A}) = \frac{\lambda(\mathcal{A})}{\lambda(\mathcal{M})} \tag{2}$$

where $\mathcal{M}$ is measurable by definition (which means $\lambda(\mathcal{M})$ is nonzero) and finite by being compact. Hence, $\mathbb{P}(x \in \mathcal{A}) = 0$ if and only if $\lambda(\mathcal{A}) = 0$.

**Theorem 1.** *Any $P$ points sampled uniformly from a compact, Riemannian manifold embedded in $\mathbb{R}^d$ are almost surely in general position.*

*Proof.* We will prove this statement in parts based on each individual requirement that guarantees points of a set $\mathcal{S}$ are in general position in $\mathbb{R}^d$:

1. We claim no $d+1$ points are colinear or coplanar. We start with $P = 3$. Using exchangeability and independence, the probability that three points $(x_1, x_2, x_3)$ are colinear can be written as

$$\mathbb{P}(x_1, x_2, x_3 \text{ are colinear}) = \int_{x_1, x_2 \in \mathcal{M}} \mathbb{P}(x_3 \in \mathcal{A} \,|\, x_1, x_2) \mathbb{P}(x_1) \mathbb{P}(x_2) \, \mathrm{d}x_1 \, \mathrm{d}x_2 \tag{3}$$

   where $\mathcal{A}$ is the intersection of the line that goes through $x_1$, $x_2$, and $\mathcal{M}$. Here, $\mathcal{A}$ is a one-dimensional subspace of $\mathcal{M}$ and thus has measure 0, which means that the integral is 0 and the probability of colinearity is 0. For $P > 3$, there is still a finite number $\binom{P}{3}$ of combinations of points to check, all of which have probability 0. Therefore, the summation of any set being colinear is 0. We do the same proof for coplanarity. We start with $P = 4$. Again, using exchangeability and independence, the probability that four points $(x_1, x_2, x_3, x_4)$ are coplanar can be written as

$$\mathbb{P}(x_1, x_2, x_3, x_4 \text{ are coplanar}) = \int_{x_1, x_2, x_3 \in \mathcal{M}} \mathbb{P}(x_4 \in \mathcal{A} \mid x_1, x_2, x_3) \mathbb{P}(x_1) \mathbb{P}(x_2) \mathbb{P}(x_3) \, \mathrm{d}x_1 \, \mathrm{d}x_2 \, \mathrm{d}x_3 \quad (4)$$

where $\mathcal{A}$ is the intersection of the plane that goes through $x_1$, $x_2$, $x_3$, and $\mathcal{M}$. Here, $\mathcal{A}$ is a two-dimensional subspace of $\mathcal{M}$ and thus has measure 0, which means that the integral is 0 and the probability of colinearity is 0. For $P > 4$, there is still a finite number $\binom{P}{4}$ of combinations of points to check, all of which have probability 0. Therefore the summation of any set being coplanar is 0.

2. We claim no $d + 2$ points are cocircular or cospherical. We can follow a similar argument as what was made in the first point except, this time, $\mathcal{A}$ is the intersection of a translation and scaling of $\mathcal{S}^1$ or $\mathcal{S}^2$ which have 0 measure in 2 and 3 dimensions, respectively. Again all combinations checked for any value of $P$ add to 0.

3. We claim no smallest circumsphere of two, three, or four points has radius equal to $\alpha$. For two points to have a circumsphere of radius $\alpha$, the second point would have to be $2\alpha$ distance away from the first. This means that the second point lands on the sphere centered at the first point with radius $2\alpha$, which has zero measure. Therefore, probability of this occurring is 0. For three points (given that two points are not distance $2\alpha$ apart), there are two possible spheres of radius $\alpha$ for the third point to land on which would lead to this degenerate case. Both have measure 0 and, therefore, this case in summation has measure 0. For four points (given the first three points which define a circle), there are exactly two spheres of radius $\alpha$ on which the fourth could land—both of which have measure 0. Therefore, the probability of this case is also 0.

4. We claim no points lie on the smallest circumsphere of two or three other points. Similar to the second proof, once we have the circumsphere defined by a first set of points, the probability that another point lands on the circumsphere is 0.

As each of the degenerate cases occurs with probability 0 then, with probability one, none of them occur; and the sampled points are almost surely in general position. □

#### 1.2 Theoretical bounds on number of sampled points needed for new shapes

Now that we have established that the degenerate cases almost surely do not come up when sampling points from Euclidean space, we now move on to the question of how many points need to be sampled. The exact number and the bounds on that number depend on the application. For example, one may want to avoid having any isolated points, as these may not be intrinsically meaningful and statistically may be interpreted as noise. In this case, one may decide to discard isolated points; however, on the other hand, some may not want to be faced with this decision at all. The following result shows that the probability of isolated points goes to zero as the number of sampled points goes to infinity. This also leads to a result which gives a minimum bound on the number of sampled points needed in or order to have no isolated points (with high confidence).

**Theorem 2.** *Let $\mathcal{M}$ be a compact submanifold, let $\mathbb{P}$ denote the probability measure over that manifold, and let $\alpha > 0$ be fixed. If $P$ points $\{x_1, ..., x_P\}$ are sampled i.i.d. according to $\mathbb{P}$, then the probability that there are no isolated points in the resulting $\alpha$-complex approaches 0 as $P \to \infty$.*

*Proof.* First, let us only consider the case were $\alpha < \text{diam}(\mathcal{M})/2$, since an $\alpha$ value out of that range gives us a convex hull of points. We can also assume that the support of $\mathbb{P}$ is over all of $\mathcal{M}$; otherwise, we can adjust our work to just be over the closure of the support of the probability measure. We will start by assuming that there is exactly one isolated point. By exchangeability (given by the independent sampling), we can consider this the first point sampled which we will denote as $x_1$. For this point to be isolated, it must be an $\alpha$-extreme point (and, therefore, not on the interior of the shape)—that is, there must be some ball of radius $\alpha$, which we will denote $\mathcal{B}_\alpha$, such that $x_1$ lies on the boundary and this ball does not contain any other sampled points. Additionally, there must be no points such that there is a ball of radius $r < \alpha$ such that $x_1$ and another point lie on the boundary of the ball. In other words, all other points must be more than a $2\alpha$ distance away. Since $x_1$ was sampled, we know that $\mathbb{P}(\mathcal{B}_{2\alpha}(x_1)) > 0$ since there must be support for the probability measure within the ball. By the definition of a probability measure, the probability that the next sampled point is outside of this ball is $\mathbb{P}(x \in \mathcal{B}^c_{2\alpha}(x_1)) < 1$. The probability that the first point sampled is isolated from the rest of the $P-1$ points can be written as

$$\begin{aligned}\mathbb{P}(x_1 \text{ is isolated}) &= \int_{\mathcal{M}} \mathbb{P}(x \in \mathcal{B}^c_{2\alpha}(x_1))^{P-1}\mathbb{P}(x_1)\,\mathrm{d}x_1 \\ &\leq \int_{\mathcal{M}} \left[1 - \min_{x_1\in\mathcal{M}} \mathbb{P}(x \in \mathcal{B}_{2\alpha}(x_1))\right]^{P-1} \mathbb{P}(x_1)\,\mathrm{d}x_1 \\ &= \left[1 - \min_{x_1\in\mathcal{M}} \mathbb{P}(\mathcal{B}_{2\alpha}(x_1)\cap\mathcal{M})\right]^{P-1} \int_{\mathcal{M}} \mathbb{P}(x_1)\,\mathrm{d}x_1 \\ &= \left[1 - \min_{x_1\in\mathcal{M}} \mathbb{P}(\mathcal{B}_{2\alpha}(x_1)\cap\mathcal{M})\right]^{P-1}.\end{aligned} \tag{5}$$

Since we have established that the minimum probability of $\mathcal{B}_{2\alpha}(x_1)$ is greater than 0, this whole expression approaches 0 as $P \to \infty$. A similar proof can be repeated for each number of finite isolated points, where instead of considering the minimum measure of a ball, we consider the minimum measure of $K$ non-overlapping balls to get the same result. Thus, the probability that any point is isolated is the sum of the probabilities that exactly $K$ points are isolated, where $1 \leq K \leq R$ with $R$ being the number of balls of size radius $2\alpha$ which fill $\mathcal{M}$, approaches 0 as $P \to \infty$. □

**Theorem 3.** *When sampling uniformly, the minimum number of points needed such that the probability of an isolated point in the $\alpha$-shape is less than some $\delta > 0$ is given by*

$$\frac{\ln(\delta)}{\ln\left[1 - \min_{x\in\mathcal{M}} \mathbb{P}(\mathcal{B}_{2\alpha}(x)\cap\mathcal{M})\right]} + 1 < P \tag{6}$$

*where* $\ln(x)$ *denotes the natural log of* $x$*.*

*Proof.* After some algebra, we know from Theorem 2 that the bound can be guaranteed if

$$\left[1 - \min_{x\in\mathcal{M}} \mathbb{P}(\mathcal{B}_{2\alpha}(x)\cap\mathcal{M})\right]^{P-1} < \delta$$

Taking natural log on both sides of this relationship gives:

$$(P-1)\ln\left[1 - \min_{x\in\mathcal{M}} \mathbb{P}(\mathcal{B}_{2\alpha}(x)\cap\mathcal{M})\right] < \ln(\delta)$$

Since $[1 - \min_{x\in\mathcal{M}} \mathbb{P}(\mathcal{B}_{2\alpha}(x)\cap\mathcal{M})] < 1$, the natural log this quantity is negative; so dividing by this number on both sides flips the inequality sign. Adding 1 to both sides then gives the desired result. □

Of course, having no isolated points with high confidence leaves room for variation in the homology of the shapes being sampled. One may not care about this; indeed, a potential extension of our work lies in purely exploring the shapes generated from a particular probability distribution. In some cases, such as in reconstructing new shapes from a data set, a user may need to preserve the homology and restrict variation to only occur between the geometric features on the boundary itself. In this case, we need a bound that can guarantee that the open ball cover will preserve the homology of the submanifold. Such a bound exists—it is calculated and proven in Niyogi et al. (2008) so long as there is a limit on the reach (which, to keep consistent with the main text, we will denote as $\tau$). Given a compact submanifold $\mathcal{M}$ embedded in $\mathbb{R}^d$, define a set $\mathcal{G}$ by the following

$$\mathcal{G} = \left\{ x \in \mathbb{R}^d \mid \exists\, p, q \in \mathcal{M} \text{ where } \operatorname{dist}(x, \mathcal{M}) = \|x - p\| = \|x - q\| \right\}$$

where $p$ and $q$ are distinct points and $\operatorname{dist}(x, \mathcal{M}) = \inf_{s \in \mathcal{M}} \|x - s\|$ is the distance from $x$ to the manifold $\mathcal{M}$. The *medial axis* of $\mathcal{M}$ is the closure of $\mathcal{G}$ and we can then define the reach by the following

$$\tau = \inf_{p \in \mathcal{M}} \sigma(p)$$

where $\sigma(p)$ is the distance of point $p$ to the medial axis. Informally, this $\tau$ is the bound of (i) the distance between multiple connected components of $\mathcal{M}$ and (ii) the radius of the smallest hole or void in $\mathcal{M}$. For a shape that is a boundary object (for example, a closed path in two dimensions or a surface mesh in three dimensions), $\tau$ is the smallest distance between a point on the shape and the medial axis. We note that, in cases where $\tau$ is infinite (e.g., a convex shape), the algorithm for estimation will still find a finite number given the simplicial complexes of the input data. With this definition we can apply the main result from Niyogi et al. (2008) to our work (slightly rewritten to match our notation):

**Theorem 4.** *Let $\mathcal{M}$ be a compact $L$-dimensional submanifold of $\mathbb{R}^d$ with reach $\tau$. Let $\mathcal{S}$ be the set of $P$ points drawn i.i.d. uniformly from $\mathcal{M}$. Let $0 < \alpha < \tau/2$, $\vartheta_1 = \arcsin(\alpha/8\tau)$, $\vartheta_2 = \arcsin(\alpha/16\tau)$, and*

$$\beta_1 = \frac{vol(\mathcal{M})}{(\cos^L(\vartheta_1))vol(\mathcal{B}^L_{\alpha/4})}, \qquad \beta_2 = \frac{vol(\mathcal{M})}{(\cos^L(\vartheta_2))vol(\mathcal{B}^L_{\alpha/8})}, \qquad \mathcal{U} = \bigcup_{p \in \mathcal{S}} \mathcal{B}_\alpha(p).$$

*Then for all $\beta_1 [\log \beta_2 - \log \delta] < P$, the homology of $\mathcal{U}$ equals the homology of $\mathcal{M}$ with high confidence (probability greater than $1 - \delta$).*

Since $\mathcal{R}_p(\alpha) = \mathcal{B}_\alpha(p) \cap \mathcal{V}(p) \subset \mathcal{B}_\alpha(p)$, it follows that $\cup_{p\in\mathcal{S}}\mathcal{R}_p(\alpha) \subset \mathcal{U}$. Since $\cup_{p\in\mathcal{S}}\mathcal{R}_p(\alpha)$ covers $\mathcal{U}$, it follows that $\mathcal{U} = \cup_{p\in\mathcal{S}}\mathcal{R}_p(\alpha)$. Note the $\alpha$-complex $\mathcal{S}_\alpha$ and $\cup_{p\in\mathcal{S}}\mathcal{R}_u(\alpha)$ are homotopy equivalent (Edelsbrunner and Harer, 2010) by the Nerve Theorem; furthermore, since each $\mathcal{R}_p(\alpha)$ is convex, and therefore contractible, $\mathcal{S}_\alpha$ has the same homology as the cover. Thus, the bound above is the same bound needed for an $\alpha$-shape to preserve the homology of the original manifold.

#### 1.3 Probabilistic models for generating $\alpha$-shapes with known manifold $\mathcal{M}$

We now state our proposed naïve probabilistic model for sampling $\alpha$-shapes by drawing random points from a compact submanifold $\mathcal{M}$ embedded in $\mathbb{R}^d$

$$x_1, \ldots, x_P \overset{i.i.d.}{\sim} \text{Unif}(\mathcal{M}), \qquad P \sim P_{min}(\alpha) + \pi(\varphi), \qquad \alpha \sim \rho(\tau) \tag{7}$$

where $x_1, \ldots, x_P$ are points in $\mathbb{R}^d$ that are i.i.d. uniformly sampled over the manifold $\mathcal{M}$; $P$ is the number of points in the $\alpha$-shape with parameter $\alpha$; $P_{min}(\cdot)$ is a function of $\alpha$ based on the desired characteristics of the new generated shape (e.g., no isolated points, preserving the homology); $\pi$ is a probability distribution with hyperparameter $\varphi$; and $\rho$ is another probability distribution that is parameterized by the reach $\tau$. There are a few important takeaways from Eq. (7). First, in general, $\rho$ only needs to be some probability distribution with support between 0 and $\tau/2$ so, in the `ashapesampler` software package, we use a truncated normal distribution such that

$$\alpha \sim TN(\mu, \nu^2, 0, \tau/2). \tag{8}$$

with mean $\mu$, variance $\nu^2$, lower bound 0, and upper bound $\tau/2$. Of course, this is not the only choice. Alternatively, one could always use a standard normal distribution and instead force $0 < \mu < \tau/2$ with sufficiently small $\nu^2$. Another option would be to make $\alpha$ a deterministic function of $\tau$ to reduce the amount of topological variation we see in generated shapes (e.g., what we do in the main text by setting $\alpha = \tau - \epsilon$). Second, the value of $P_{min}(\alpha)$ will be determined by the desired level of variation between the newly generated shapes. If one wants to sample the space freely, then $P_{min}$ should be set to 0. If one does not want any isolated points, then $P$ is dictated by a user-chosen probability parameter $\delta$ and the minimum volume of an intersection of a ball of radius $\alpha$ and the manifold $\mathcal{M}$ (see next section for details).

If preserving the homology of the manifold, then $P_{min}(\alpha)$ is dictated by $\delta$, $\alpha$, and the reach $\tau$. The third important takeaway from Eq (7) is that the probability $\pi$ is an arbitrary discrete distribution with support on the natural numbers and some hyperparameter $\varphi$. This can be ignored if one does not want variation in the number of points across generated shapes. Finally, while we sample points uniformly from the manifold itself, other distributions could in theory be used so long as they have support on the manifold. Note, however, that moving away from a uniform does change the proof for the bound preserving the homology of the manifold in Theorem 4.

### 2 Avoiding Isolated Points when Generating 2D Shapes

In this section, we show how to practically calculate the minimum number of points needed such that the probability of an isolated point in a generated 2D $\alpha$-shape is less than some $\delta > 0$ (Theorem 3). Here, we need to find the minimum possible overlap between a ball of radius $2\alpha$ and the manifold that we are sampling points (i.e, $\mathcal{B}_{2\alpha}(x) \cap \mathcal{M}$). Below, we will say that the ball of interest has radius $\tilde{\alpha} = 2\alpha$ to simplify notation.

#### 2.1 Area of overlap between a circle and square

Consider a square with a side of length $r$ and a circle (or ball) of radius $\tilde{\alpha}$. When the circle is on the boundary of the square, the smallest overlap occurs when the circle is centered at one of the square's four corners. There are three cases to consider when calculating the area of the intersection between the circle and square:

(I) $0 < \tilde{\alpha} < r$;

(II) $r < \tilde{\alpha} < r\sqrt{2}$;

(III) $r\sqrt{2} \leq \tilde{\alpha}$.

In the first scenario, the area of the overlap is simply a quarter of the area of the circle: $A = \pi\tilde{\alpha}^2/4$. In the third scenario, the radius of the circle is bigger than the diagonal of the square and, thus, the circle covers the square. The area of the intersection, in this case, is the area of the square itself: $A = r^2$.

The second scenario is the one that requires more care. When $r < \tilde{\alpha} < r\sqrt{2}$, the area of intersection is equal to two half-segments subtracted from the quarter of the circle (see Fig S6 for an example). By symmetry, this means that we are effectively calculating the segment formed by a chord distance $r$ from the center of the circle. Using trigonometry, the angle formed by the two line segments of radius $\tilde{\alpha}$ from the center of the circle to the endpoints of the chord is given by $\vartheta = 2\arccos(r/\tilde{\alpha})$. We can then plug this into the area of a segment $a = [\vartheta - \sin(\vartheta)]\alpha^2/2$. Thus, the area of overlap in this scenario is $A = \pi\tilde{\alpha}^2/4 - a$. Note that we do not have to consider any special angles for $\arccos(\cdot)$ because, as we can see in Fig S6, $\vartheta/2 < \pi/4$ for all cases in the second scenario.

#### 2.2 Area of overlap between two circles

Let $\tilde{\alpha}$ denote the radius of a circle centered around a point $p$ on the circumference of the disk of radius $r$. There are three cases to consider when calculating the area of overlap between two circles where one is centered on the circumference of the other:

(I) $0 < \tilde{\alpha} < r\sqrt{2}$;

(II) $r\sqrt{2} \leq \tilde{\alpha} < 2r$;

(III) $2r \leq \tilde{\alpha}$.

In the first scenario, we can compute the area of overlap $A$ by calculating each respective circular segment, which we denote $A_1$ and $A_2$. Using the law of cosines, we find $\vartheta_1 = 2\arccos(\tilde{\alpha}^2/2r\tilde{\alpha})$ to be the central angle of the circle of radius $r$ and $\vartheta_2 = 2\arccos(2r^2 - \tilde{\alpha}^2/2r^2)$ to be the central angle of the circle of radius $\tilde{\alpha}$. Then the circular segment for each is given by the following, respectively,

$$A_1 = \frac{1}{2}[\vartheta_1 - \sin(\vartheta_1)]\tilde{\alpha}^2, \qquad A_2 = \frac{1}{2}[\vartheta_2 - \sin(\vartheta_2)]r^2. \tag{9}$$

As a result, we just need $A = A_1 + A_2$ to get the total area (see Fig S7 for an illustration). The second scenario is a little more complicated since, once $\tilde{\alpha} > r\sqrt{2}$, the angle $\vartheta_2$ becomes larger than $\pi$. While the formulas stay the same for calculating $\vartheta_1$, $\vartheta_2$, and $A_1$, the formula for $A_2$ needs to be adjusted to

$$A_2 = \frac{1}{2}[\vartheta_2 + \sin(2\pi - \vartheta_2)]r^2 \tag{10}$$

One can derive the formula in Eq (10) by seeing that we are now adding a triangle to a segment rather than subtracting one from a segment. In the third scenario, the circle of radius $\tilde{\alpha}$ completely covers the disk and so the area of intersection is simply the area of the disk: $A = \pi r^2$.

##### 2.3 Area of overlap between an annulus and circle

Consider the scenario where a circle with radius $\tilde{\alpha}$ is centered on the major border of an annulus which has major (or outer) radius $R$ and minor (or inner) radius $r$. The scenarios to consider when calculating the area of overlap are the following:

(I) $0 < \tilde{\alpha} < R - r$;

(II) $R - r < \tilde{\alpha} < R + r$;

(III) $R + r < \tilde{\alpha} < 2R$;

(IV) $2R < \tilde{\alpha}$.

For the first scenario, we can just use the same formula for the overlapping circles from the previous section. In the second scenario, we are (i) calculating the overlap of circles with radii $R$ and $\tilde{\alpha}$, where the second circle is centered on the circumference of the first, and then (ii) subtracting the overlap between the circles with radii $\tilde{\alpha}$ and $r$. The former can be calculated using the same methodology for the overlapping circles as described in the previous subsection. The latter, however, has a slightly different angle calculation as illustrated in Fig S8 but finding the overlap is straightforward once the angle is calculated. For the third scenario, we have covered the entire inner hole of the annulus, so the area of the overlap is major radius $R$ minus $\pi r^2$. In the fourth scenario, we have completely covered the annulus so the total area of the overlap is $\pi(R^2 - r^2)$. Finally, while there are the other scenarios with larger angles to be considered as before, we choose not go into detail as their solutions end up being similar to the overlapping circles cases in the previous section.

#### 3 Avoiding Isolated Points when Generating 3D Shapes

In this section, we show how to practically calculate the minimum number of points needed such that the probability of an isolated point in a generated 3D $\alpha$-shape is less than some $\delta > 0$ (Theorem 3). As was done in the 2D shape case, we need to find the minimum possible overlap between a ball of radius $2\alpha$ and the manifold that we are sampling points (i.e, $\mathcal{B}_{2\alpha}(x) \cap \mathcal{M}$). Again, we will continue to say that the ball of interest has radius $\tilde{\alpha} = 2\alpha$ to simplify notation.

#### 3.1 Volume of overlap between a cube and sphere

Consider a cube with a side of length $r$ and a sphere of radius $\tilde{\alpha}$. When the sphere is on the boundary of the cube, the smallest overlap occurs when the cube is centered at one of the cube's eight corners. There are four cases to consider when calculating the area of the intersection between the cube and sphere:

(I) $0 < \tilde{\alpha} \leq r$;

(II) $r < \tilde{\alpha} \leq r\sqrt{2}$;

(III) $r\sqrt{2} < \tilde{\alpha} < r\sqrt{3}$;

(IV) $r\sqrt{3} \leq \tilde{\alpha}$.

The first and fourth scenarios are relatively straightforward. In the first scenario, the volume of the overlap between the shapes is an eighth of the volume of the sphere: $V = \pi\tilde{\alpha}^3/6$. In the fourth scenario, the ball covers the cube completely, so the volume of intersection is $V = 4\pi r^3/3$. In the second scenario, where $r < \tilde{\alpha} \leq r\sqrt{2}$, the sphere extends beyond the cube but does not cover the three faces which abut the corner on which the sphere is centered (see Fig S9 for an example). This creates three distinct quarter-sphere "caps" (or the three-dimensional equivalent of segments) which are part of the eighth of sphere being considered but fall outside the intersecting space. The formula for a spherical cap is

$$V_c = \frac{1}{3}\pi h^2(3u - h) \tag{11}$$

where $u$ is the radius of a general sphere and $h$ is the height of the cap. Translating the above to the second scenario in our case, $u = \tilde{\alpha}$ and $h = \tilde{\alpha} - r$. We want to consider a quarter of the cap three times and subtract that from the eighth of sphere being considered, which gives us the following volume

$$\begin{aligned} V &= \frac{1}{6}\pi\tilde{\alpha}^3 - 3\left(\frac{V_c}{4}\right) \\ &= \frac{1}{6}\pi\tilde{\alpha}^3 - \frac{1}{4}\pi(\tilde{\alpha} - r)^2(2\tilde{\alpha} - r). \end{aligned}$$

In the third scenario, where $r\sqrt{2} < \alpha < r\sqrt{3}$, the spherical caps which were considered in the second scenario now overlap. This means that we need to add back the volume from the intersection of the caps themselves, else we end up subtracting parts of the sphere twice. Let $a$ denote the base radius of the spherical cap in question, which we can calculate as $a = \sqrt{\tilde{\alpha}^2 - r^2}$. Then we have the base given by radius $a$ and the part that needs isolating has an extra length $a - r$. Using the Pythagorean theorem, the height is given by $b = \sqrt{a^2 - r^2}$ (see Fig S10 for an illustration). If we assume that the sphere of radius $\tilde{\alpha}$ is centered at the origin in three-dimensional space then, to get the volume, we just need to integrate $f(x, y) = \sqrt{\tilde{\alpha}^2 - x^2 - y^2}$ over $[0, b]$ on the x-axis and $[r, a]$ on the y-axis. Formally, the volume that needs to be subtracted is given by

$$V_o = \int_a^r \int_0^b \sqrt{\tilde{\alpha}^2 - x^2 - y^2} \, \mathrm{d}x \, \mathrm{d}y. \tag{12}$$

By symmetry, we need to subtract three of these volumes from three partial sphere caps (again with $u = \tilde{\alpha}$ and $h = \tilde{\alpha} - r$) to get the total volume to subtract from an eighth of the sphere volume. Combining this information, we get the following

$$V = \frac{1}{6}\pi\tilde{\alpha}^3 - 3\left[\frac{1}{4}V_c - V_o\right]$$

which completes these cases.

#### 3.2 Volume of overlap between two spheres

Similar to Section 2.2, we want to find the area of overlap between two spheres where one is centered on the surface of the other. This effectively comes down to finding calculating the spherical caps of the overlap between a sphere with radius $r$ with another sphere with radius $\tilde{\alpha}$ (see Fig S11). Let $V_1$ and $V_2$ denote the spherical caps for each sphere, respectively. We can use similar formulas as in the two-dimensional setting to find the polar angles, with $\vartheta_1 = \arccos(\tilde{\alpha}^2/2r\tilde{\alpha})$ being the central polar angle of the sphere with radius $r$ and $\vartheta_2 = \arccos(2r^2 - \tilde{\alpha}^2/2r^2)$ being the central polar angle of the sphere with radius $\tilde{\alpha}$. Next, we use $\vartheta_1$ and trigonometry to get the height of the spherical cap for $V_2$: $h_2 = \tilde{\alpha}\cos(\vartheta_1)$. Finally, we get $h_1 = \tilde{\alpha} - h_2$. We can now calculate both of the spherical caps as follows:

$$V_1 = \frac{1}{3}\pi h_1^2(3\tilde{\alpha} - h_1), \qquad V_2 = \frac{1}{3}\pi h_2^2(3r - h_2) \tag{13}$$

The total volume of overlap is given by $V = V_1 + V_2$. Note that when $h_2 > r$, we have to adjust the formula for $V_2$. In this situation, we now have $h_* = 2r - h_2$ and $V_2^* = 4\pi r^3/3 - \pi h_*^2(3r - h_*)/3$. This scenario is illustrated in Fig S12. Furthermore, in the special case where $2r < \tilde{\alpha}$, the volume of the overlap is just volume of the sphere with radius $r$.

##### 3.3 Volume of overlap between a shell and sphere

In this section, we now want to compute the volume of overlap when a sphere of radius $\tilde{\alpha}$ is centered at the boundary of a shell. Here, a shell refers to the volume between two concentric spheres, one of radius $R$ and the other of radius $r$ (where $R > r$ without loss of generality). The three scenarios to consider are:

1. $0 < \tilde{\alpha} < R - r$
2. $R - r < \tilde{\alpha} < R + r$
3. $R + r < \tilde{\alpha} < 2R$
4. $2R < \tilde{\alpha}$

For three of these scenarios, we can follow the same logic we detailed in the previous section when considering overlapping spheres. In the first scenario, the intersecting volume is just the overlap between the sphere with radius $R$ and the sphere of radius $\tilde{\alpha}$. It is the same overlap in the third scenario, but we just need to subtract the volume of the smaller sphere with radius $r$. In the final scenario, the volume over overlap is the volume of the shell. In the second scenario, we are (i) taking the overlap between the spheres with radii $\tilde{\alpha}$ and $r$, and (ii) subtracting the overlap between the spheres with radii $\tilde{\alpha}$ and $R$. For the smaller sphere, we can again use the law of cosines on the triangles with sides $\tilde{\alpha}$, $r$, and $R$ to calculate the angles we need. The calculation of $h_2 = \tilde{\alpha}\cos(\vartheta) - R$ is the adjusted for the shell, as well as $h_1 = \tilde{\alpha} - R - h_2$. Afterwards, all subsequent calculations follow the same logic as the previous section.

#### 4 Uniformly Sampling Points from a Known Manifold

In this section, we detail how to uniformly sample new points when the true underlying manifold $\mathcal{M}$ is known (see Eq (7)). Here, we focus on a disk in two-dimensions and a solid ball in three-dimensions. Extensions to the annulus and shell, respectively, can be easily derived. While sampling from the unit square or cube is relatively straightforward, more care needs to be taken when sampling from a disk or ball. This is because, while the parameters used are polar coordinates defined over clear bounds, these do not necessarily have to follow uniform distributions. Indeed, rejection sampling can be used to get uniformity in both cases; however, we can use alternative methods with better efficiency.

#### 4.1 Sampling uniformly from a 2D disk

In this section, we will use polar coordinates $(r, \vartheta)$ and convert them back into Cartesian coordinates $(x, y)$ since that is usually the form in which we will receive shape data. Let $b$ be the fixed radius of a ball such that $r \in [0, b]$ and $\vartheta \in [0, 2\pi]$. Typically, we can convert a polar coordinate pair $(r, \vartheta)$ to Cartesian coordinates using the following formula

$$x = r\cos(\vartheta), \qquad y = r\sin(\vartheta). \tag{14}$$

To convert Cartesian coordinates to polar coordinates, we can use the following

$$r = \sqrt{x^2 + y^2}, \qquad \vartheta = \operatorname{sign}(y)\arccos(x/r). \tag{15}$$

The area of a disk is $A = \pi b^2$, so the uniform probability density function (pdf) for $(x, y)$ is given by

$$f(x, y) = \frac{1}{\pi b^2}\mathbb{I}_{\{x^2+y^2\leq b^2\}}(x, y) \tag{16}$$

where $\mathbb{I}(\cdot)$ is an indicator function. To get the probability density for $(r, \vartheta)$, we use change of variables. After some algebra, we get that the determinant of the Jacobian matrix for $\mathrm{d}(x, y)/\,\mathrm{d}(r, \vartheta)\,(r, \vartheta)$ is $r$. Then, after substituting in Eq (15) into the indicator function, we get the pdf for $(r, \theta)$ as

$$f(r, \vartheta) = \frac{r}{\pi b^2}\mathbb{I}_{\{0\leq r\leq b\}}(r)\mathbb{I}_{\{0\leq\vartheta\leq 2\pi\}}(\vartheta). \tag{17}$$

While $\vartheta$ is still uniform over $[0, 2\pi]$, the marginal distribution for $r$ is $f(r) = 2r/b^2$ which has a cumulative distribution function (CDF) of the form $F(r) = r^2/b^2$ and thus is not uniform over $[0, b]$. To remedy this, we use inverse transform sampling to estimate $r$. Extension to the annulus just requires that we change the lower bound of the range of $r$ to some nonzero number instead of 0.

#### 4.2 Sampling uniformly from a 3D ball

We can use the same change of variables strategy to obtain a cumulative distribution function (CDF) for three-dimensional polar coordinates $(r, \vartheta, \phi)$, where now $\vartheta$ is the horizontal angle and $\phi$ is the vertical angle. Let $\phi \in [0, \pi]$ and, once again, let $r \in [0, b]$ and $\vartheta \in [0, 2\pi]$. Given a set of polar coordinates, we can convert to cartesian coordinates using the following formula

$$x = r\cos(\vartheta)\sin(\phi), \qquad y = r\sin(\vartheta)\sin(\phi) \qquad z = r\cos(\phi). \tag{18}$$

Similarly, to convert Cartesian coordinates to polar coordinates, we can use the following

$$r = \sqrt{x^2 + y^2 + z^2}, \qquad \vartheta = \arccos\left(\frac{x}{\sqrt{x^2 + y^2}}\right), \qquad \phi = \arccos(z/r). \tag{19}$$

Since the volume of a solid ball in three-dimensions is $V = 4\pi b^3/3$, the probability density function (pdf) for the cartesian coordinates is given by the following

$$f(x, y, z) = \frac{3}{4\pi b^3}\mathbb{I}_{\{x^2+y^2+z^2<b^2\}}(x, y, z). \tag{20}$$

After substituting Eq 19 into the indicator variables and computing the Jacobian matrix determinant to be $r^2\sin(\phi)$, we find that the pdf in polar coordinates is

$$f(r, \vartheta, \phi) = \frac{3r^2\sin(\phi)}{4\pi b^3}\mathbb{I}_{0\le r\le b}(r)\mathbb{I}_{0\le\vartheta\le 2\pi}(\vartheta)\mathbb{I}_{0\le\phi\le\pi}(\phi) \tag{21}$$

Once again, $\vartheta$ remains uniform over $[0, 2\pi]$, but $r$ and $\phi$ are not uniform over their respective ranges. Indeed, the CDF for $r$ is $F(r) = r^3/2b^3$, and the CDF for $\phi$ is $F(\phi) = -[\cos(\phi) - 1]/2$. For both of these variables, we use inverse transform sampling to simulate uniformity. Extensions to the shell would again require changing the lower bound of $r$ to some nonzero number.

### 5 Supplementary Figures

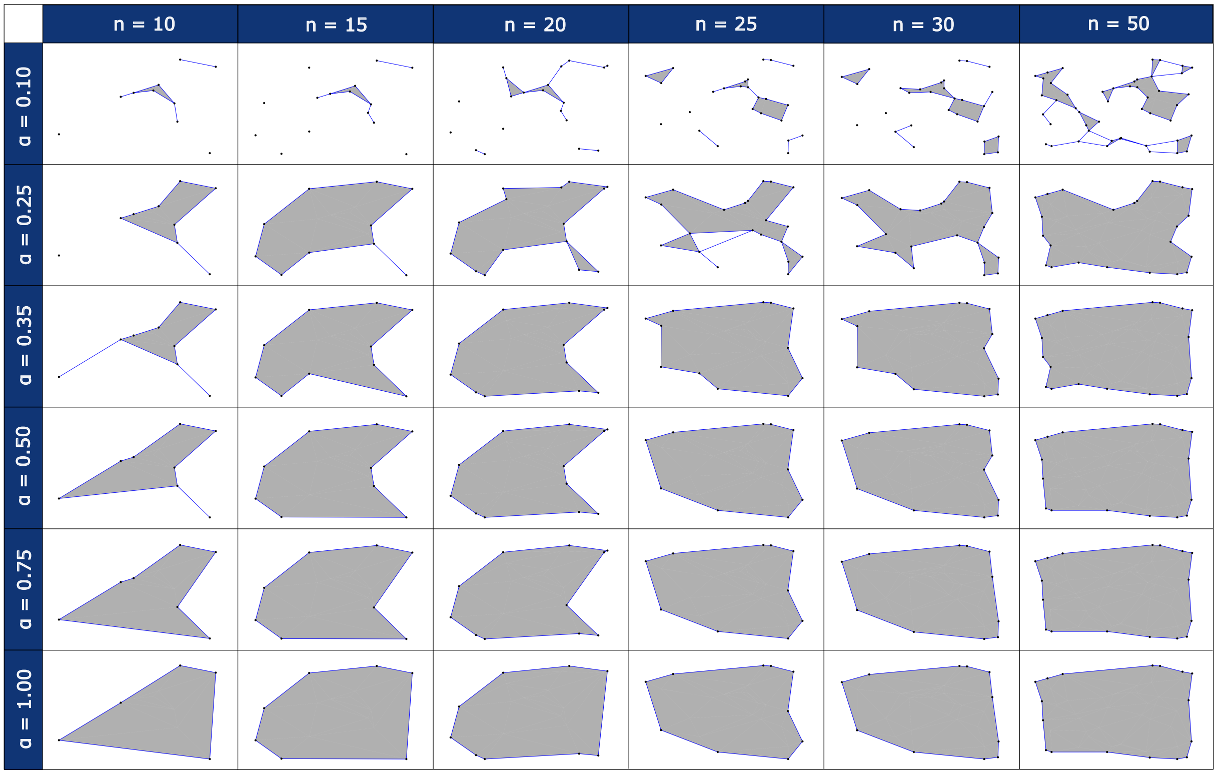

**Figure S1. Visual table showing $\alpha$-shapes being generated as a function of the number of points $P$ sampled from a unit square and the parameter $\alpha$.** Each column uses the same $P$ points for consistency and ease of comparison across rows. Additionally, the points used in a given column are a subset of the points used in columns to the right.The first row has the smallest $\alpha$ value and many disconnected points; while, the final row has one solid connected shape with no holes. Parameters $P$ and $\alpha$ need to be chosen based on (i) how much detail is desired, (ii) what characteristics in the local and global structure we want to preserve, and (iii) how close to the original manifold we want for the generated $\alpha$-shape.

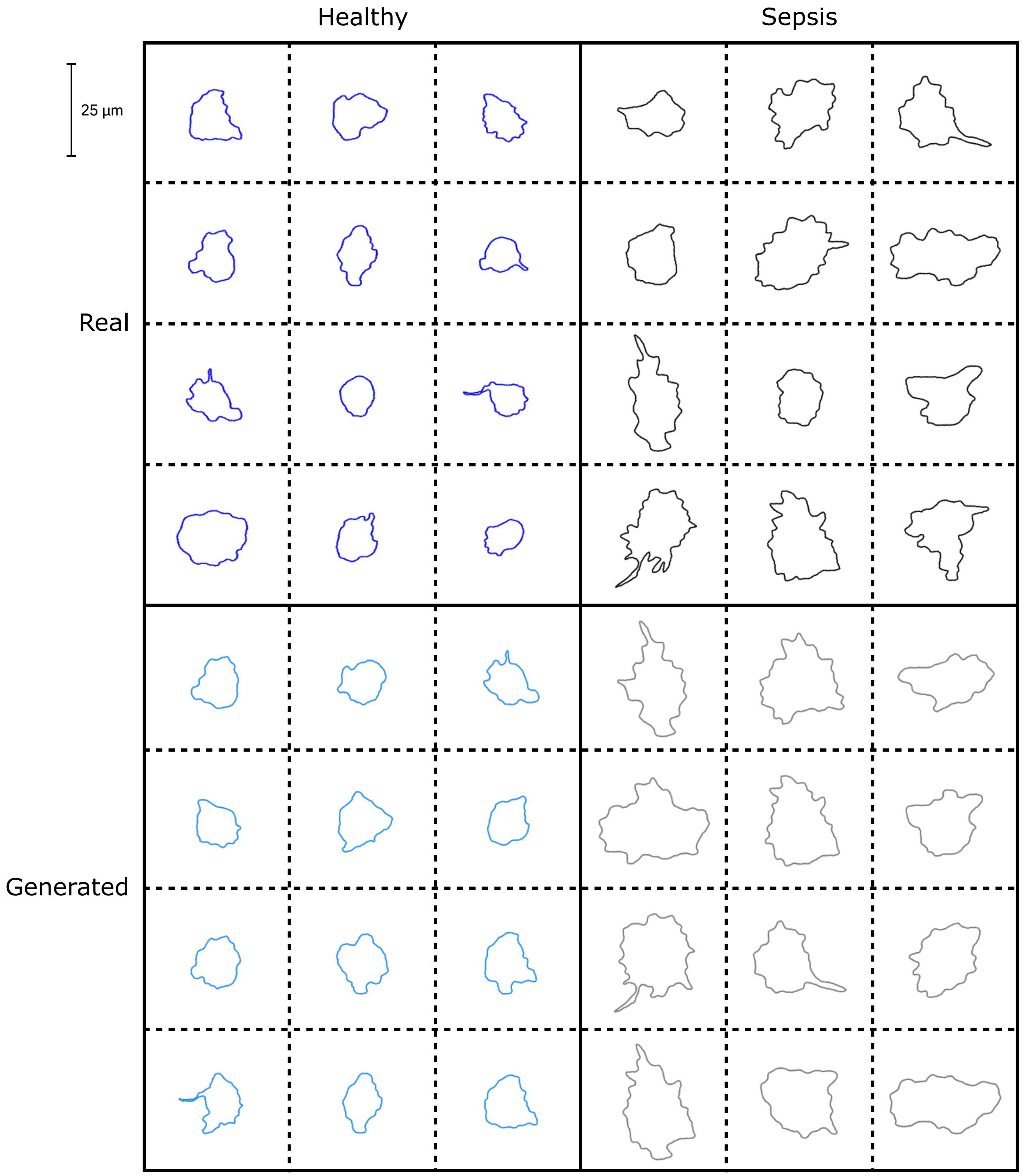

**Figure S2. Additional samples of real healthy (blue), generated healthy (light blue), real septic (black), and generated septic (gray) neutrophils in gels with stiffness 1.5 kilopascals (kPa).** Immediately noticeable are the differences in area and number of protrusions between the healthy versus septic neutrophils. With the additional neutrophils in each category, it also becomes clear that there is greater heterogeneity in the septic neutrophil class as compared to the healthy neutrophils.

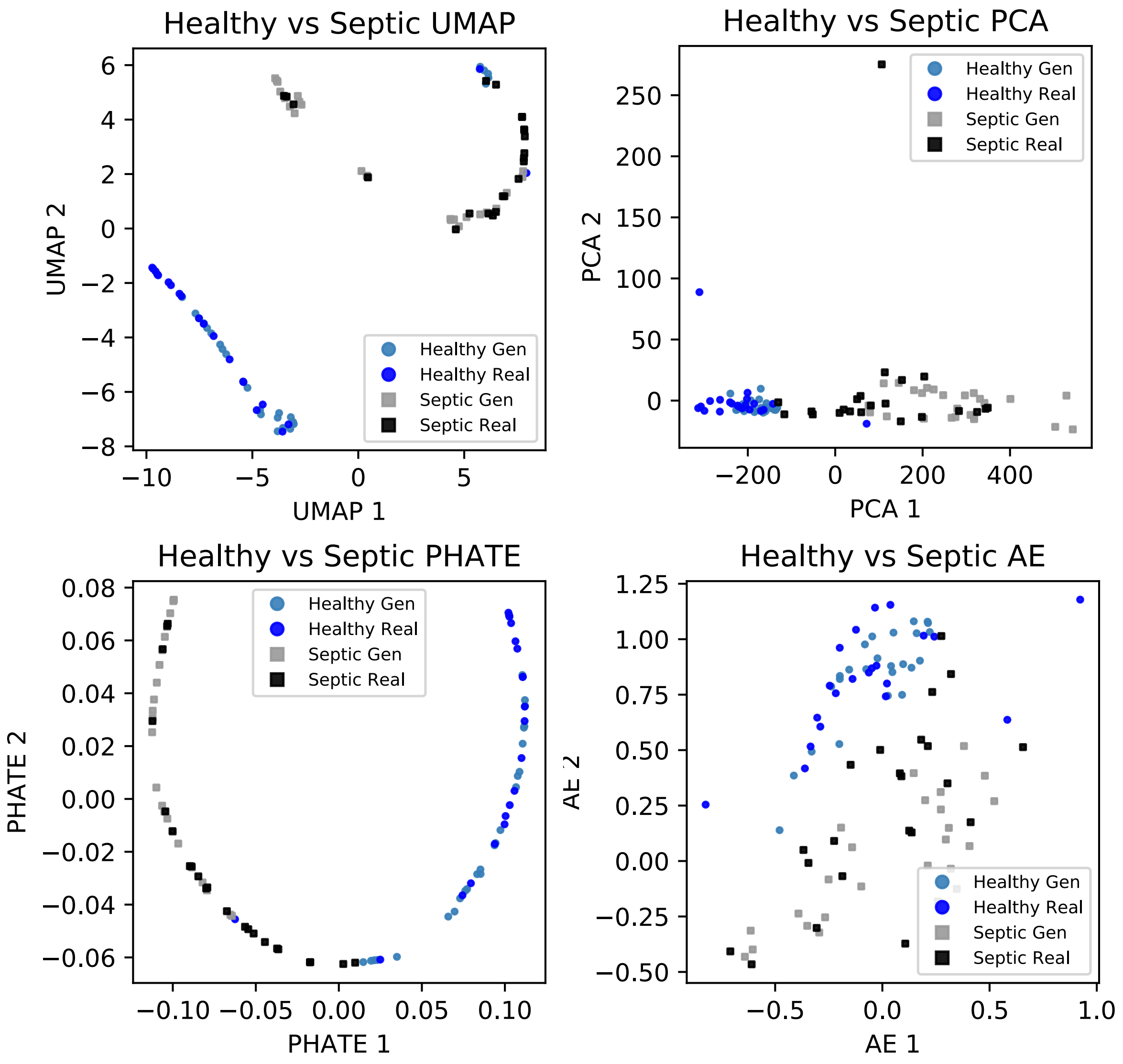

**Figure S3. Additional dimensionality reduction plots showing clusters of real and generated healthy and septic neutrophils.** Other dimensionality reduction approaches including: the uniform manifold approximation projection (UMAP), PHATE, principal component analysis (PCA), and a generic autoencoder. In all plots, the neutrophil types form distinct clusters, and the generated cells from the $\alpha$-shape sampler are mixed in among the corresponding neutrophil class from which they were referenced.

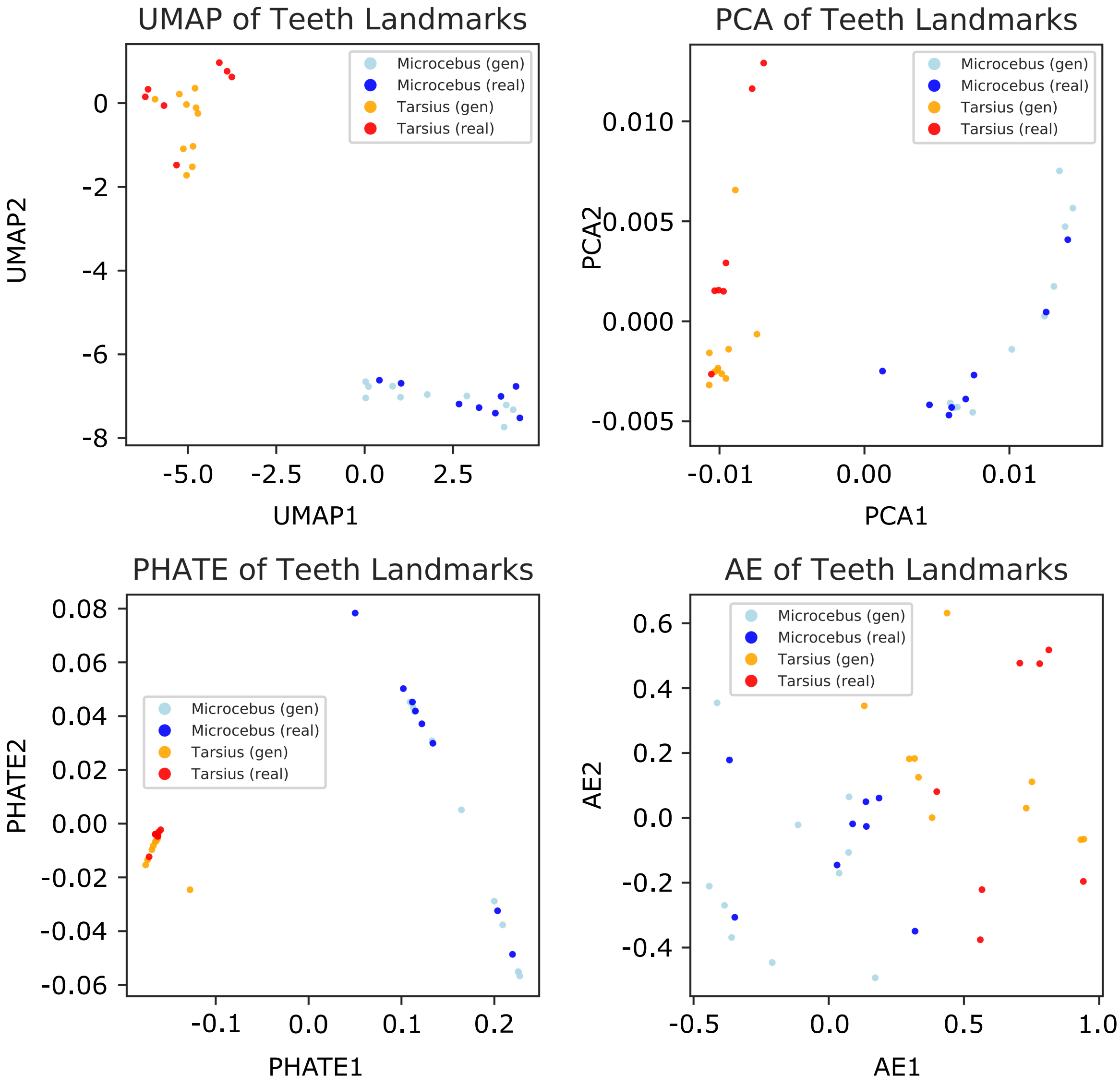

**Figure S4. Additional dimensionality reduction plots showing clusters of real and generated *Microcebus* and *Tarsius* teeth.** Other dimensionality reduction approaches including: the uniform manifold approximation projection (UMAP), PHATE, principal component analysis (PCA), and a generic autoencoder. In all plots, the two species form distinct clusters, and the generated molars from the $\alpha$-shape sampler are mixed in among the corresponding primate genus from which they were referenced.

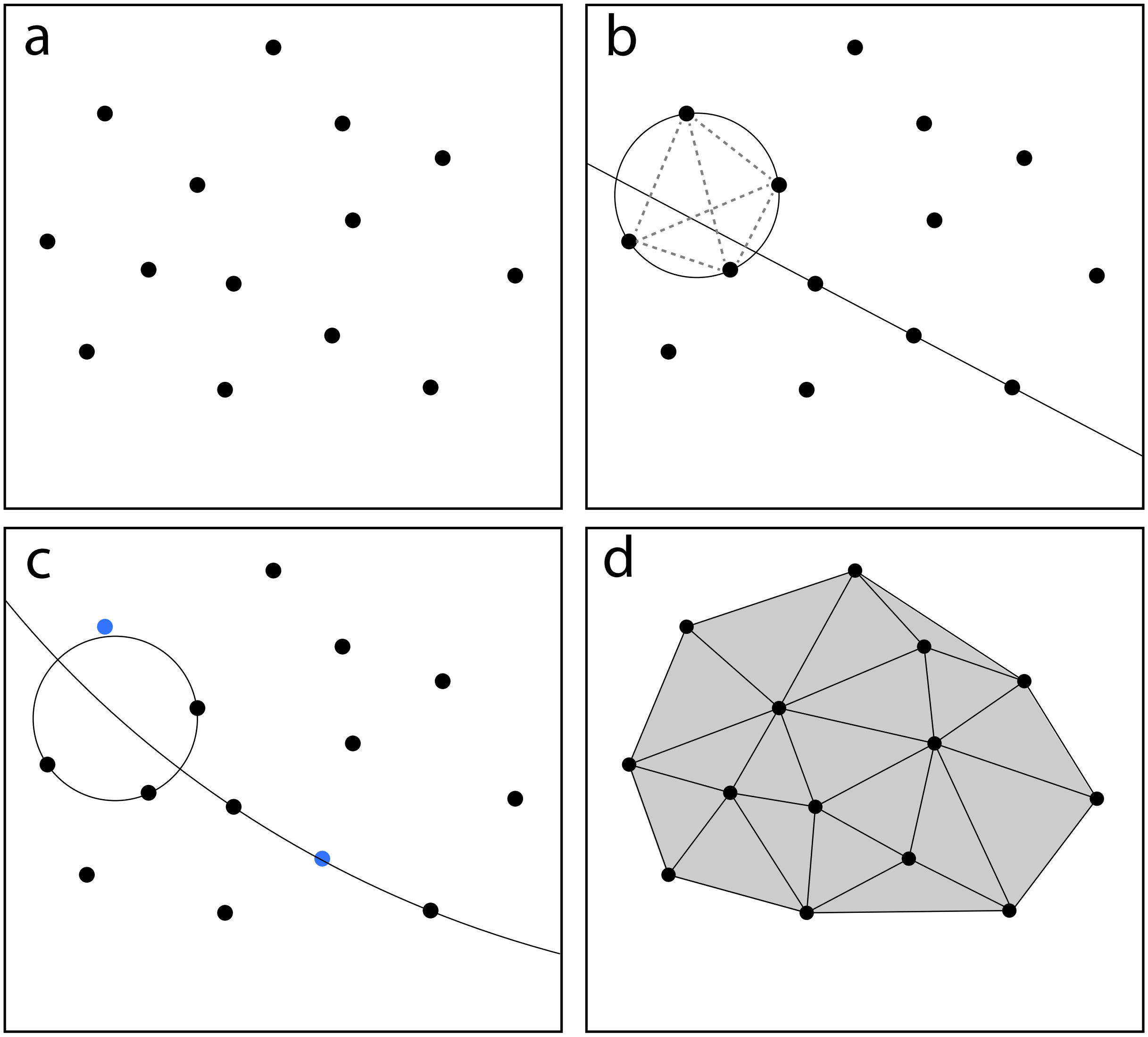

**Figure S5. Two-dimensional illustration of the importance of points being in general position (i.e., no three points colinear, no four points cocircular) iin order to be able to create an $\alpha$-complex, which is a subcomplex of the Delaunay complex.** In panel **(a)**, consider a set of points in two-dimensions. Delaunay triangulation checks all possible sets of three points and the unique circles they form. If there are no other points from the point set within the circle, those three points form a face (otherwise they do not). Panel **(b)** shows the issues that can arise when points are not in general position. If three points are colinear, then there is no circle to check. Regardless of whether or not this is an issue from an algorithmic perspective, topologically, there are now multiple features representing what could be one edge. When four points are cocircular, there are now multiple ways to connect the points and uniqueness is lost. In panel **(c)**, we show that we can perturb some or all points to solve for this—but then there is an issue of which points to perturb and this additional strategic decision adds an extra layer to the generative pipeline which we want to avoid if possible. Panel **(d)** shows the Deluanay triangulation of the points given in panel **(c)**. Any $\alpha$-complex on these points will be a subcomplex of this Delaunay complex.

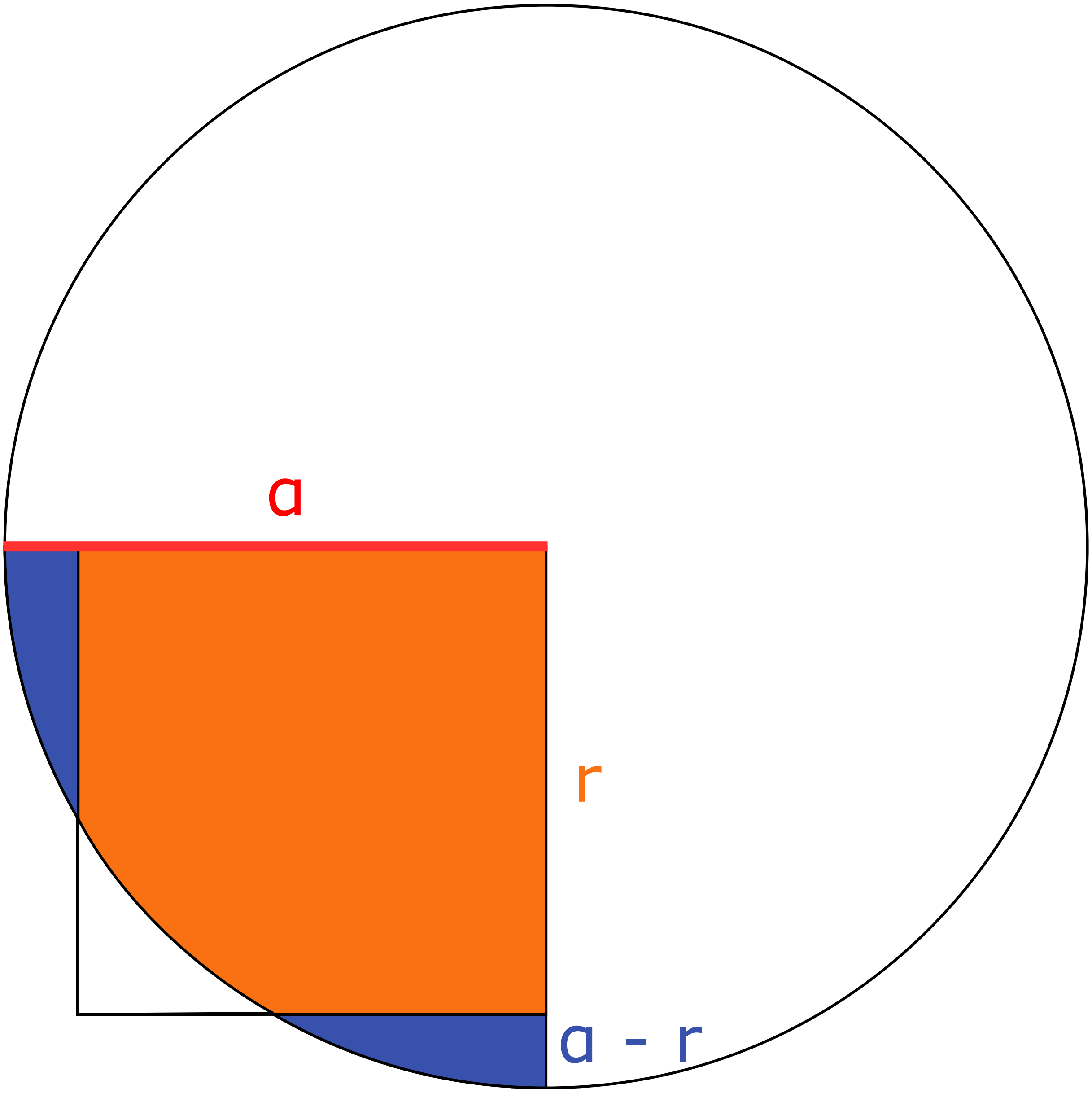

**Figure S6. Diagram illustrating the overlapping area of a circle with radius $\tilde{\alpha}$ and a square with side length $r$ when $r < \tilde{\alpha} < r\sqrt{2}$ and the circle is centered on the corner of the square.** The shaded areas make up a quarter of the circle: the orange portion is the area of interest and the blue region is the part of the circle we need to subtract. Using symmetry, we see that the total blue area is the segment of height $\tilde{\alpha} - r$, so that area is what we subtract from the quarter circle to get the orange area of interest.

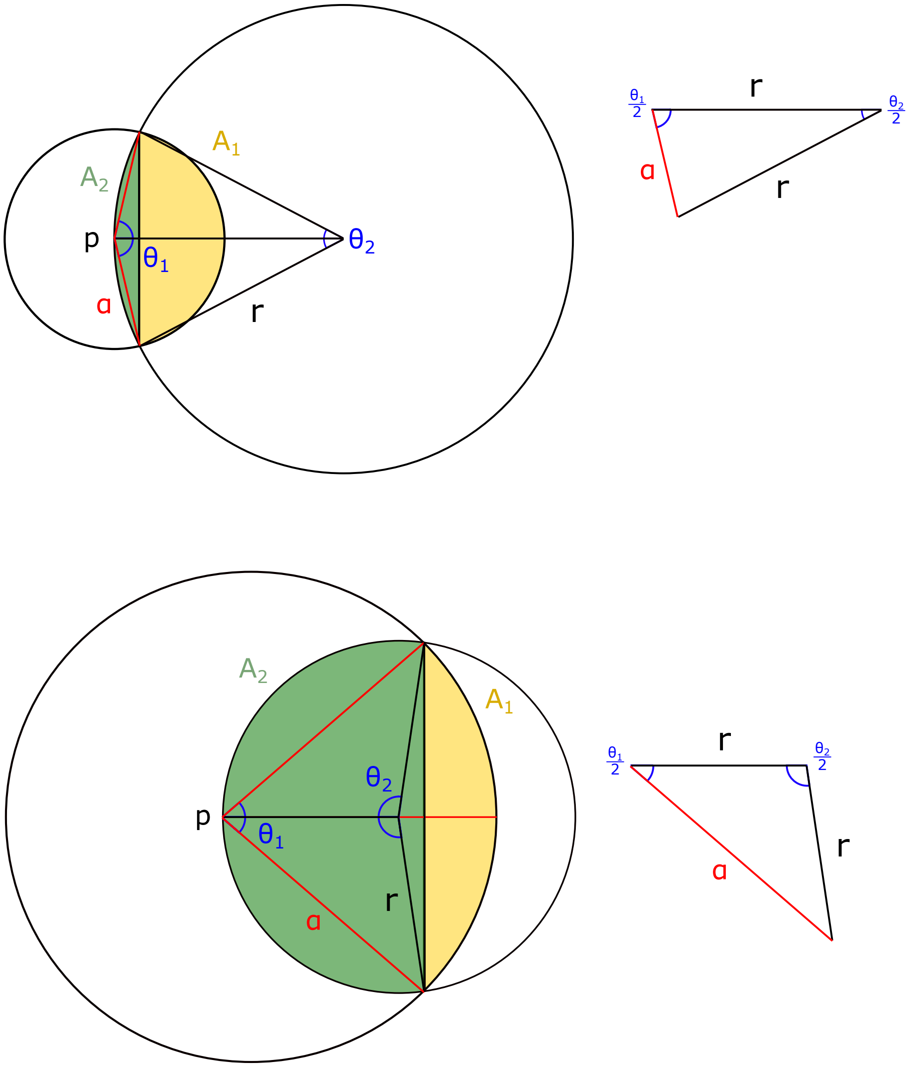

**Figure S7. Diagram illustrating two overlapping circles where one circle is centered at the circumference of the other.** One circle has a center at say point $p$ with radius $\tilde{\alpha}$, while the other circle has radius $r$. We need to use the law of cosines to find $\vartheta_1$ and $\vartheta_2$ so that we may calculate $A_1$ and $A_2$. The top half of the figure is the scenario where $0 < \tilde{\alpha} < r\sqrt{2}$, while the bottom half is the scenario where $r\sqrt{2} < \tilde{\alpha} < 2r$.

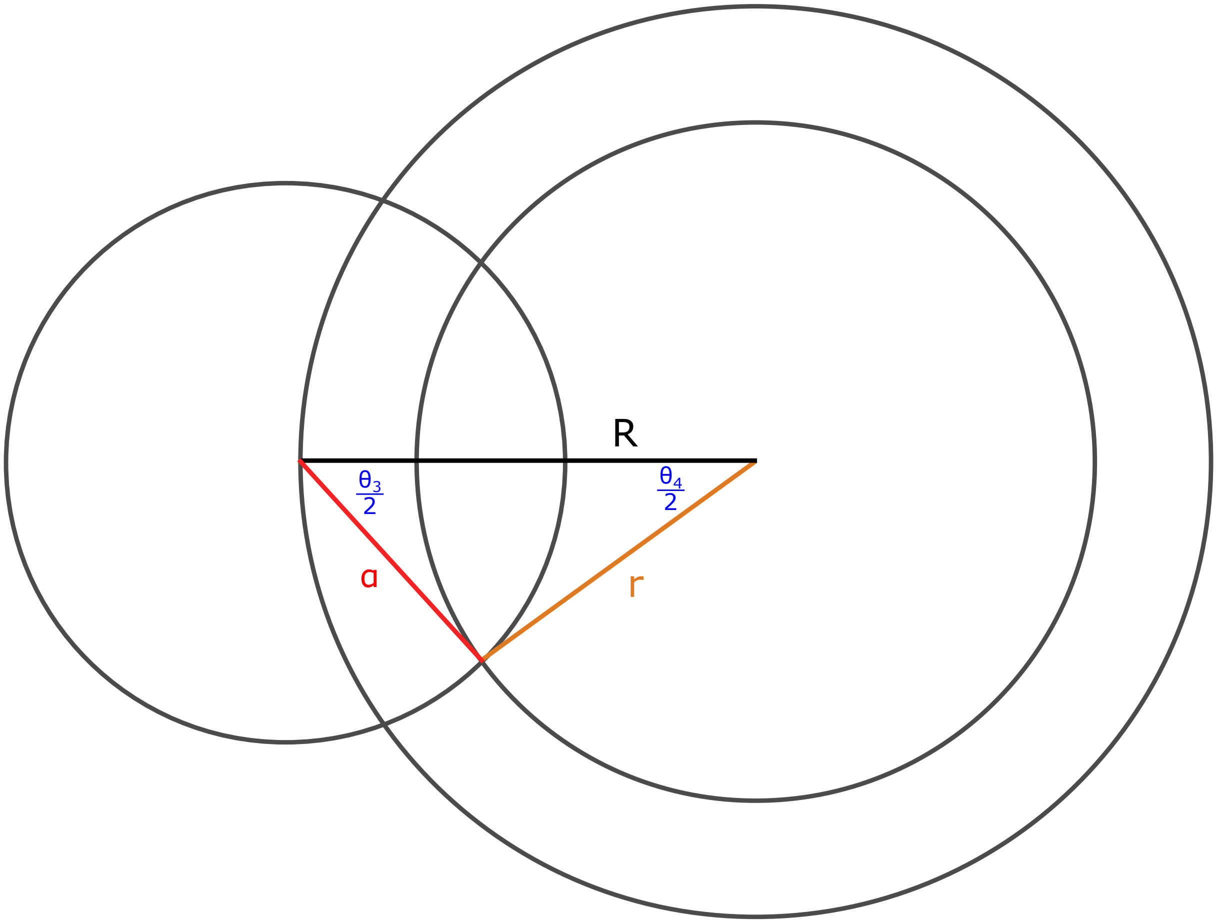

**Figure S8. Diagram illustrating the overlap between a circle of radius $\tilde{\alpha}$ and an annulus with major radius $R$ and minor radius $r$.** Here, the circle with radius $\tilde{\alpha}$ is centered on the circumference of the circle with radius $R$. In particular, this is the scenario where $R - r < \tilde{\alpha} < R + r$. We need to use a slightly different triangle to use the law of cosines to calculate $\vartheta_3$, and we will need to use $\vartheta_4$ to calculate the segments making up the overlap with the hole of the annulus. Note that we have enough information to find these parameters.

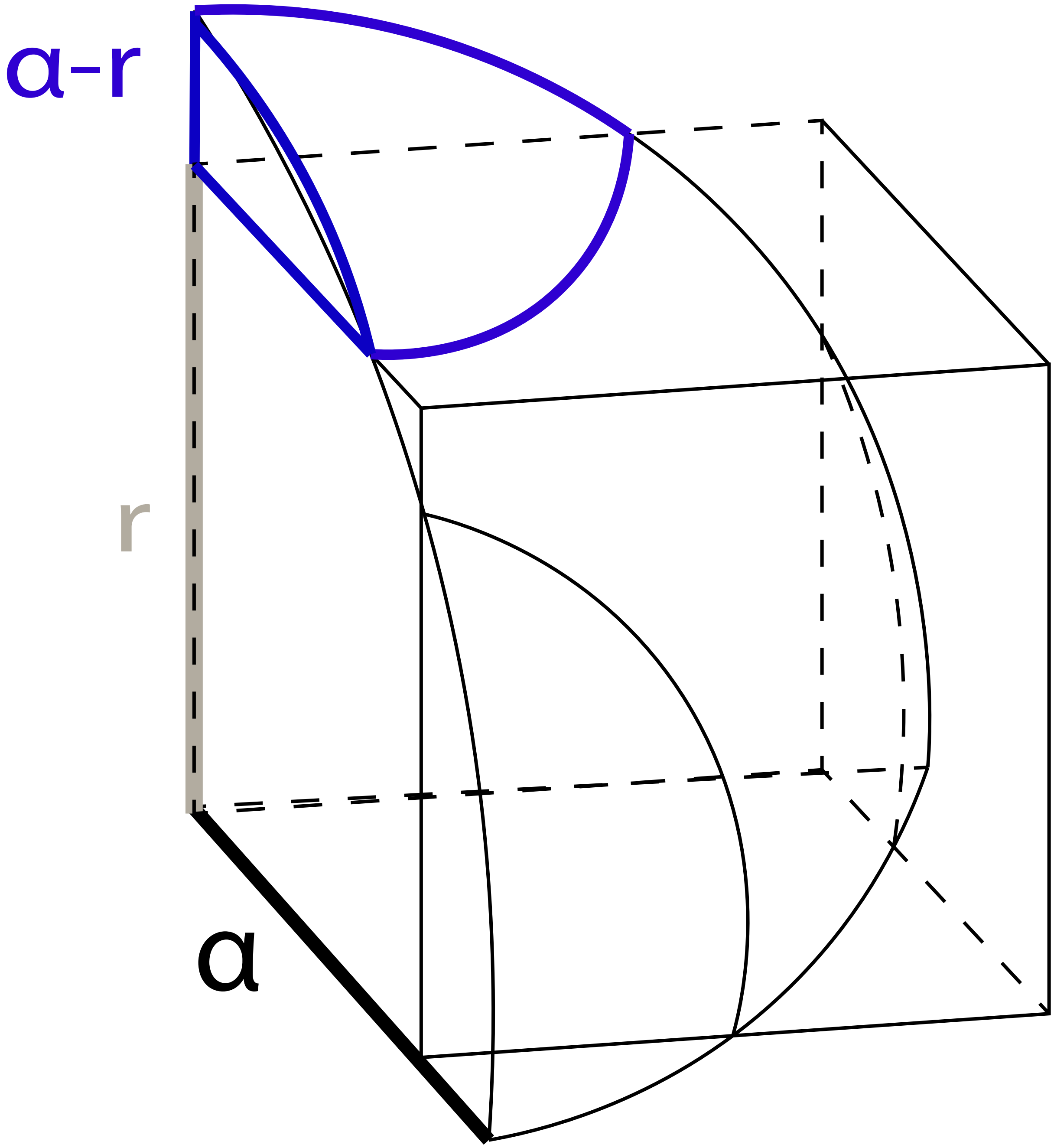

**Figure S9. Illustration where one eighth of a sphere with radius $\tilde{\alpha}$ is overlapping a cube with side $r$, where $r < \tilde{\alpha} < r\sqrt{2}$.** To get the volume of overlap between a sphere and a cube, we only need to consider one eighth of the sphere. To do so, we calculate three "extra" protruding volumes (one of which is highlighted in blue) which we then subtract from the total sphere volume. These extra volumes are all equal by symmetry and we can obtain them by using the formula for a spherical cap.

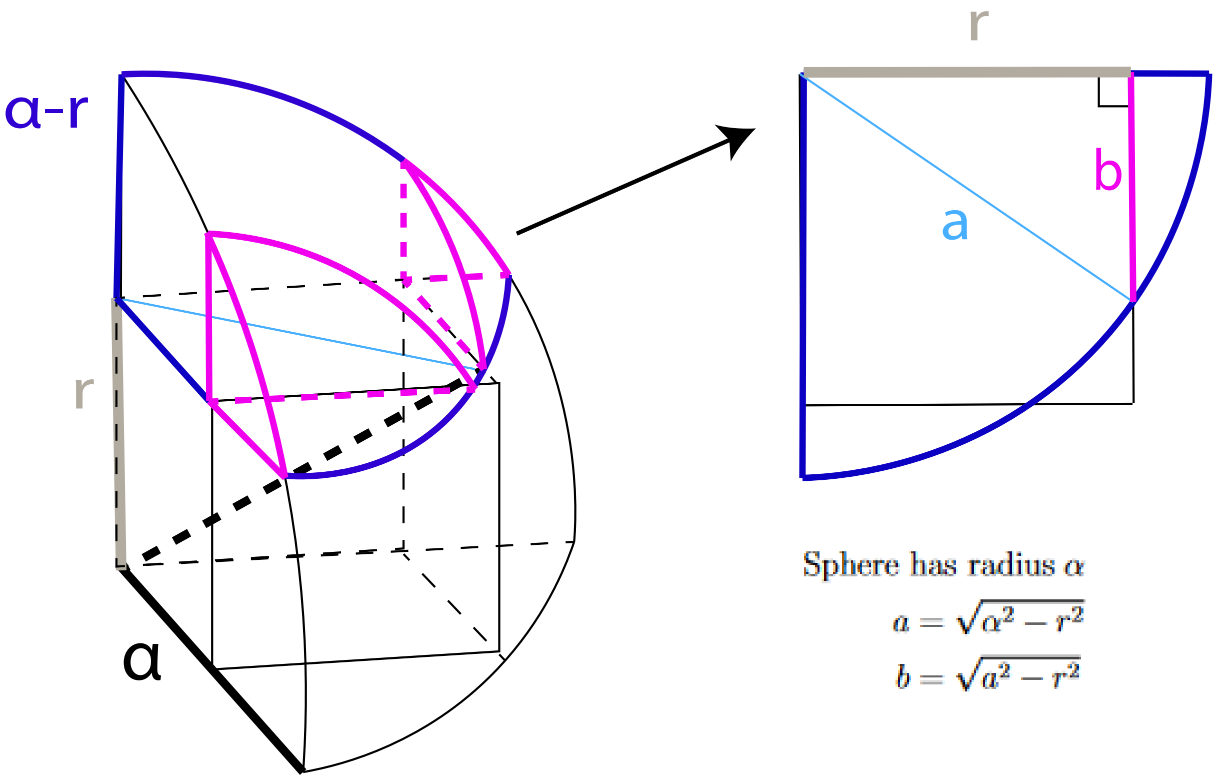

**Figure S10. Illustration of a case where a sphere with radius $\tilde{\alpha}$ is overlapping a cube with side $r$, where $r\sqrt{2} < \tilde{\alpha} < r\sqrt{3}$.** This special scenario requires additional work to get the correct volume of overlap. The three extra volumes shown in Fig S9 are now overlapping, so we need to add back information from these intersections. We can again do this by using the spherical cap formula, but we need to make sure we get the correct radius for that cap. We can use the pythaogrean theorem to get the value of $a$, which is the distance from the corner of the cube to the sphere along the top face of the cube. We can then take that slice and use the Pythagorean Theorem again to obtain the correct value of $b$.

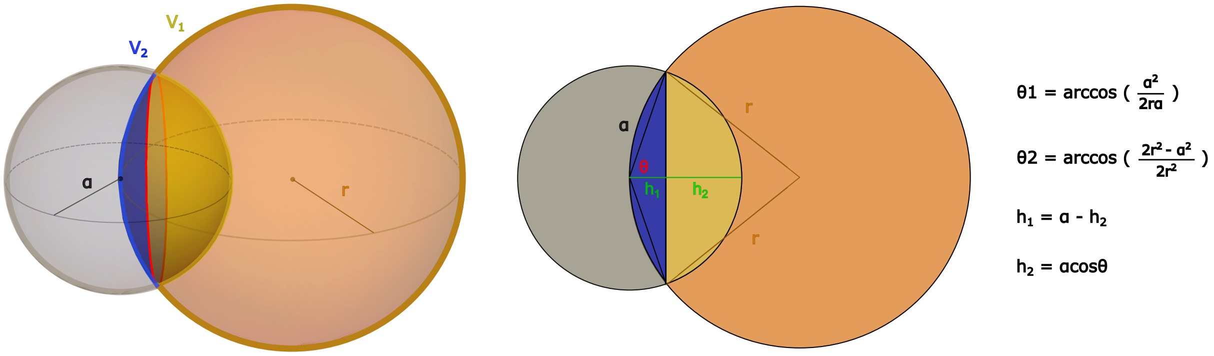

**Figure S11. Illustration of the scenario where two spheres overlapping, both from a three-dimensional perspective (left) and a two-dimensional perspective (right).** Calculating the intersection comes down to calculating the volume of two spherical caps and adding those volumes together. We can use trigonometry to calculate the correct values of the heights and radii of the spherical caps.

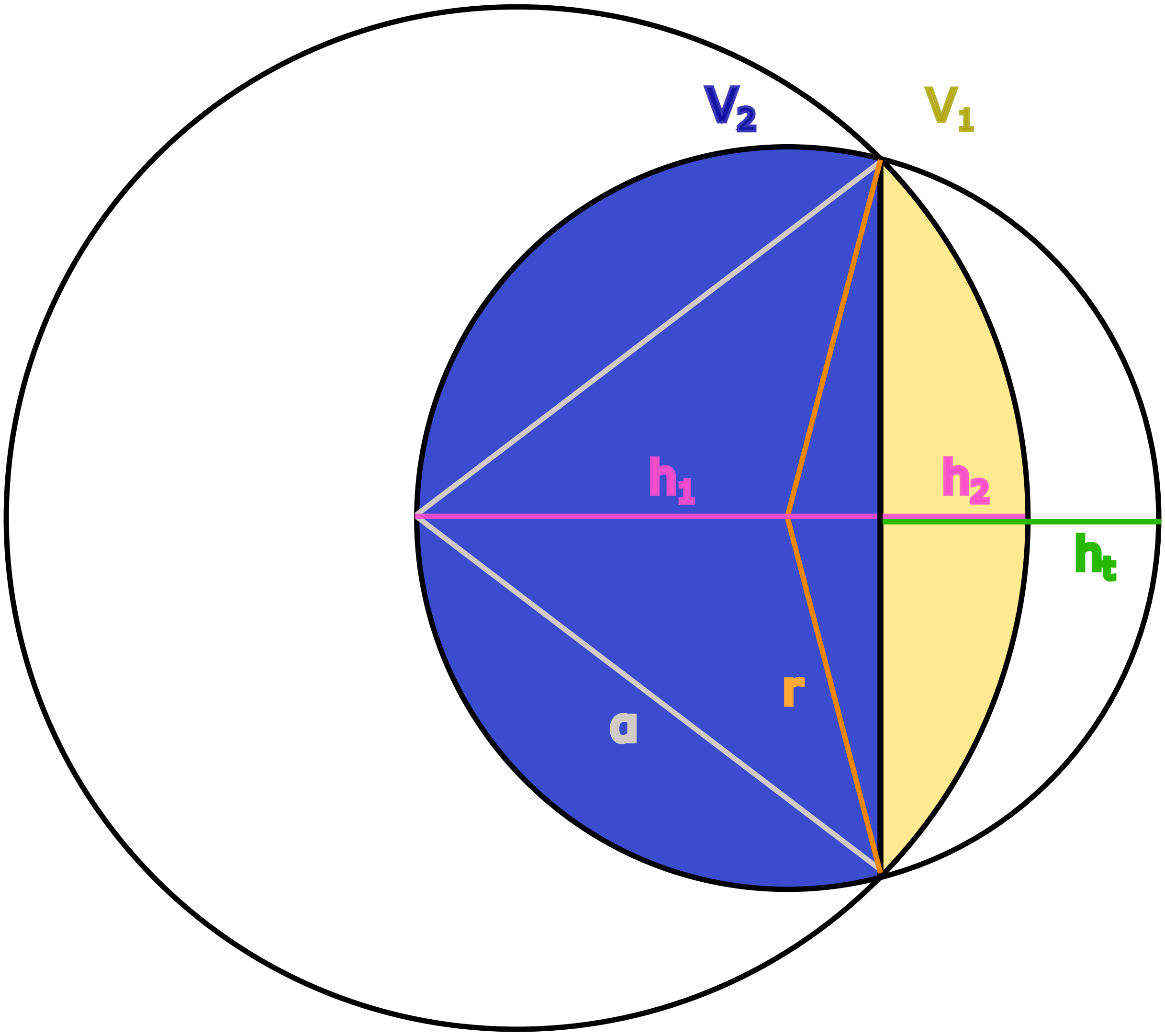

**Figure S12. Two-dimensional view of two overlapping spheres, but this time where $\tilde{\alpha} > r$.** The geometry is a little different here than in Fig S11, but the volume of the overlap is still calculable.

419 # 6 Supplementary Tables

**Table S1. Table of the inner radius, outer radius, and thickness measurement for the $N^* = 10$ generated annuli using the $\alpha$-shape sampler.** Reference annuli were created with inner radius $r = 0.25$, outer radius $R = 0.75$, and thickness equal to $R - r = 0.5$. We consider these measurements to be the "ground truth" during evaluation. The root mean square error (RMSE) is taken by comparing the original reference annuli to the generated shapes. (XLSX)

**Table S2. Table of the major and minor radii for the $N^* = 10$ generated tori using the $\alpha$-shape sampler.** Reference annuli were created with major radius $R = 0.75$ and minor radius $r = 0.25$. We consider these measurements to be the "ground truth" during evaluation. The root mean square error (RMSE) is taken by comparing the original reference tori to the generated shapes. (XLSX)

**Table S3. Comparing differences between the real and generated healthy and septic neutrophils for 33 geometric shape characteristics.** Here, we compare the differences between groups using a t-test. The corresponding test statistic and resulting $P$-value are reported in the last two columns. (XLSX)

**Table S4. Mean Euclidean distances comparing different combinations of real and generated *Microcebus* and *Tarsius* teeth.** Overall, we observe that the generated *Microcebus* and generated *Tarsius* teeth are nearly twice as close to their respective real groups than to any other group. (XLSX)
